# Interspecies relationships of wild amoebae and bacteria with *C. elegans* create environments propitious for multigenerational diapause

**DOI:** 10.1101/2024.06.03.597131

**Authors:** Marcela Serey, Esteban Retamales, Gabriel Ibañez, Gonzalo Riadi, Patricio Orio, Juan Pablo Castillo, Andrea Calixto

**Affiliations:** Centro Interdisciplinario de Neurociencia de Valparaíso, Instituto de Neurociencia, Facultad de Ciencias, Universidad de Valparaíso; Universidad de Valparaíso; ANID – Millennium Science Initiative Program Millennium Nucleus of Ion Channels-Associated Diseases (MiNICAD); Center for Bioinformatics, Simulation and Modeling, CBSM, Department of Bioinformatics, Faculty of Engineering, University of Talca. Campus Talca, Chile

**Keywords:** natural environments, amoebae, microbiota, behavior, diapause, multigenerational, RNAi, interspecies communication

## Abstract

The molecular and physical communication within the microworld supports the entire web of life as we know it. How organisms such as bacteria, amoeba and nematodes -all superabundant-interact to sustain their niche, however, is not known, especially how their associations generate and affect behavior of animals in fluctuating environments. To have a frame to study interactions between microbe and animal, we collected soil from a temperate semi-arid climate and isolated the culturable genus of bacteria *Comamonas, Stenotrophomonas, Chryseobacterium* and *Rhodococcus* and the amoeba *Tetramitus*. This ensemble was then fed in long-term experiments to the nematode *C. elegans* to study developmental rate, diapause entry, fertility, feeding behavior and neuronal integrity. We observed that the ensemble is long lasting and induces animals to diapause after a few generations under conditions that are not canonically pathogenic. We called this phenomenon Dauer Formation in Natural Ensembles (DaFNE). DaFNE requires the communication between live bacteria and the nematode intestine, suggesting the existence of a bidirectional interaction in the holobiont. While all bacteria from the ensemble colonize the intestine of the nematodes, *Comamonas* is the most represented and *Rhodococcus* the scarcest. The amoeba *Tetramitus* can be ingested by *C. elegans*, but it is not part of its microbiota.

DaFNE depends on pheromone and nematode quorum, but high temperature in the homeostatic range, triggers diapause with fewer numbers. DaFNE increases as generations pass and is also remembered transgenerationally. The RNA interference (RNAi) pathway is needed for initiation of DaFNE, indicating the communication via RNA is crucial to execute bacterially induced behaviors in natural environments.

**Significance:** Microbes have an overwhelming influence over the animals they live with, modulating development and decision making. Microscopic nematodes are the most abundant multicellular animals in the biosphere, suggesting they possess well-rehearsed successful relationships with their associated microbiota. Little is known about the modulation of nematode behavior in complex ecosystems with multiple organisms interacting. We use bacteria and amoeba from a natural ecosystem and introduce the pioneer nematode *C. elegans* to study behavioral parameters in long lasting experiments. The most striking response of nematodes to this natural environment is the commitment to diapause of a significant portion of the population. We call this form of hibernation Dauer Formation in Natural Ensembles or DaFNE. We propose that animals in nature may hibernate frequently, as a result of the communication with their natural biota. We find that DaFNE requires pheromone production in nematodes and also the RNA interference pathway, suggesting the RNA repertoire of both entities may be at play.

Higher temperatures in the optimal range for nematode growth, require much less nematode quorum for DaFNE, indicating that a non-noxious increase in temperature favors diapause in natural environments. Nematodes respond to each bacterium in different ways when grown in monocultures and in the ensemble. This suggests that the abundance of specific species in nature may shift behavioral preferences and outputs in microscopic animals. We also show that the amoeba *Tetramitus* can be ingested by worms, demonstrating that *C. elegans* is a broader microbivore. Like worms, amoebae display specific responses to bacteria and add variability to behaviors elicited by nematodes. Finally, bacteria in the ensemble unlike in monocultures, are not exhausted during the length of the experiments even in the presence of bacterivore nematodes and amoebae.

## Introduction

The relationship between ubiquitous microscopic organisms such as protists, bacteria and nematodes is essential to ecosystem health and biosphere sustainability (1–4). How these organisms interact to sustain their niche is largely understudied, in part given the myriad of variables at play in nature. A key aspect of niche health is the interplay within the microfauna to promote behavioral adaptation to superabundant nematodes. While certain aspects of the wild nematode’s biology have been explored in the context of natural bacteria (5–10), amoebae have been for the most part ignored in studies of natural interactions of nematodes and bacteria (11).

*Caenorhabditis elegans* is an excellent representative species of the larger Nematoda class to study behavioral responses to biotic variation (12–16). One of the crucial responses of nematodes to environmental change is their ability to enter quiescence (17). In the laboratory, *C. elegans* diapauses forming the dauer larvae upon crowding, food scarcity, temperature increase and bacterial pathogenesis (18–21). In natural settings, dauers are common but the cues that trigger dauer entry in this context are unknown (22, 23). Here we find that an ensemble of *Comamonas*, *Rhodococcus, Chryseobacterium* and *Stenotrophomonas* and the amoebae *Tetramitus* induces *C. elegans* to diapause without promoting stereotyped pathogenesis (24) (25). In the ensemble neither species exhausts in long term experiments suggesting Dauer Formation on Natural Ensembles (DaFNE) contributes to regulating sustainability of the ecosystem.

Adaptive responses of *C. elegans* to bacteria that cause pathogenesis bacteria (21, 26) use effectors of the RNA interference machinery at different cellular and mechanistic levels. RNAi can be transmitted to the offspring (27–29) and also produce transgenerational epigenetic inheritance (30, 31). We show that mutations in *sid-1,* essential for systemic RNAi (32); *rde-1*, needed for processing of exogenous dsRNA (33, 34), and *znfx-1,* needed for heritability of RNAi (31), are required for DaFNE, revealing a larger role for RNAi in interspecies interactions. Coherent with heritable effects of RNAi effectors, DaFNE increases intergenerationally and is also remembered transgenerationally.

## Results

### A natural microbial ensemble of bacteria and amoebae to study long term relationships with *C. elegans*

To establish a framework of various organisms interacting, we collected a soil sample from a natural urban ecosystem and identified the culturable species of amoebae and bacteria. As a representing species of the superabundant nematodes (4) we chose the pioneer organism *C. elegans,* which is ideal to study behavioral outputs in response to biotic environments and their molecular mechanisms (5, 10).

The soil sample contained one type of amoeba and diverse culturable bacteria (**Fig. 1A** and **Movie S1**). We isolated bacteria by successive passages in solid Luria Bertani (LB) media whereas amoebae grew better on Nematode Growth Media (NGM) plates with their symbiotic bacteria. Wild nematodes were also found but were not cultivated. On LB plates, bacteria separated in two subcultures, of which only one contained amoebae (**Fig. 1B**) suggesting they have specific associations with the soil microbiota.

**Figure 1.**
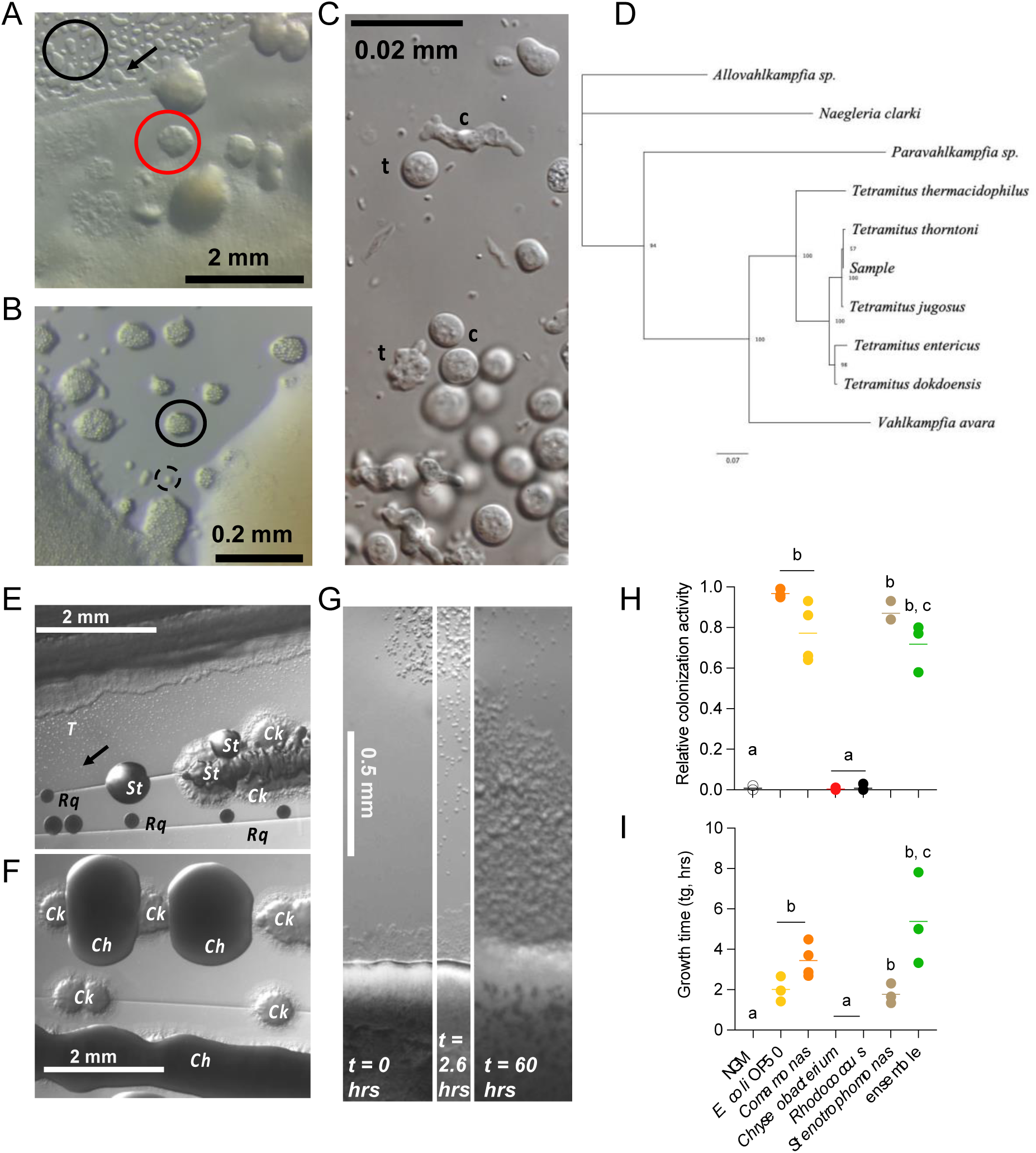
Identification of species in the ensemble. **A**. Natural microbial assembly growing on agar-NGM medium, showing distinct features. Sharp-contrast irregular structures on the agar correspond to vast patches of amoebae cysts (black circle). Round high contrast objects (red circle) are colony-like pools of bacteria and amoebae trophozoites displaying high activity as detected with time-lapse video photography (**Movie S1**). One individual cyst is marked with a black arrow. **B**. Natural microbial assembly growing on a LB plate. Note examples of patches of *Tetramitus* cysts (circle) and moving trophozoite (dashed circle). *Tetramitus* tends to accumulate on specific areas of the bacterial assembly (left), while avoiding other sectors (right). These two sectors separated spontaneously and continued to grow for several days after inoculation. **C.** DIC microphotography of cysts and trophozoites of *Tetramitus* found the original soil sample. **D.** Maximum Likelihood consensus phylogenetic inference for the sample and 5 related *Tetramitus* species, using ribosomal RNA sequence 18S and 5.8S including ITS1 and ITS2 rRNAs. The node labels show the bootstrap values based on 1,000 iterations. The tree was rooted in *Allovahlkampfia* sp. Additional species *Naegleria clarki*, *Paravahlkampfia* sp. and *Vahlkampfia avara* were used as outgroups. **E**. Colony isolation from natural soil sample on LB media, containing *Rhodococcus* (*Rq*), *Stenotrophomonas* (*St*), *Comamonas* (*Ck*) and *Tetramitus* cysts (*T*). **F**. Colony isolation from natural soil sample, containing *Comamonas* (*Ck*) and *Chryseobacterium* (*Ch*). **G**. Time-lapse experiment of *Tetramitus* amoebae colonizing the bacterial ensemble lawn. Cysts appear on top of the panel and bacterial ensemble on the bottom. placed (top of the panel) at t = 0 hrs (left). After 2.6 hrs (center) some amoebae have undergone excystation and explore the substrate reaching the bacterial lawn. 60 hours post-inoculum (right) cysts have colonized the bacterial lawn leaving cysts and trophozoites all over the recorded area. **H-I**. relative colonization activity (**H**) and growth time (**I**) in different bacterial lawns, derived from analysis of cumulative activity over time (**Fig. S2**). Experiments that do not support colonization display colonization fraction tending to zero and growth time tending to infinity (>>100 hours). Full parametric description is found in **Table 2**. Same letters denote there are no statistical differences, different letters mean statistical differences. Complete statistical analyses are detailed in **Dataset 2**.

Bacteria were identified by full length ribosomal 16S sequencing as gram-negative species of the genus *Comamonas* (**Fig. S1A)**; *Stenotrophomonas,* (**Fig. S1B**); and *Chryseobacterium,* (**Fig. S1C**); and the gram positive *Rhodococcus,* (**Fig. S1D**). Here we will refer to bacteria by their genus names (see extended Results in the Methods section for more details).

We found the amoebae in two states: cysts, which are immobile, and trophozoites that move freely on the substrate (**Fig. 1C**). We classified the amoebae isolates as a *Tetramitus* by 18S and internal transcribed spacer (ITS) sequencing. The final tree shown (**Fig. 1D**) was arbitrarily chosen among all resulting tree searches since all have the same topology and similar support values. This tree is compatible with previous trees published (35–37). The sample is shown to be related with *T. thorntoni* and *T. jugosus*, according to this phylogenetic tree which was reconstructed from 18S and ITS sequences. Trees using only ITS sequences (218 informative positions) or only 18S (1,227 informative positions) for phylogenetic reconstruction, show a similar relationship. These trees suggest that both *Tetramitus* and the sample species are closely related (see extended Results in Methods). The support values in the tree inform us that it is not possible to discriminate either species, hence we will name the amoeba by the genus name. The species forming the ensemble are described in **Table 1**.

**Table 1.**
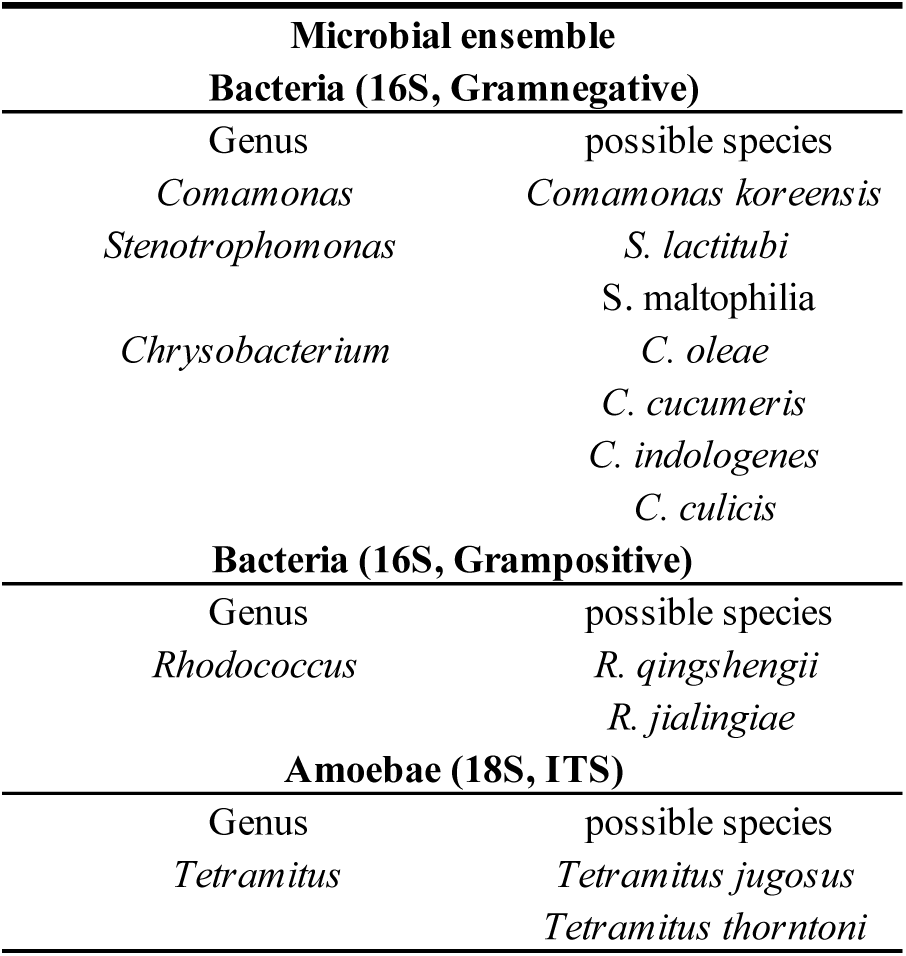
Identification of ensemble.

*Tetramitus* was found in ensembles that contained *Comamonas, Stenotrophomonas*, and to a smaller extent, *Rhodococcus* (**Fig. 1E**) and not on the mix containing *Chryseobacterium* (**Fig. 1F**). This indicates that either *Tetramitus* chooses certain bacteria or that they are more supportive of their growth. We exposed inoculums of *Tetramitus* cysts to the ensemble, monocultures and control *E. coli* OP50 and recorded their migration and growth activity patterns using time-lapse acquisition (see Methods, **Movie S2**). Cyst inoculums were placed approximately 1 mm away from the bacterial lawn and we recorded the movement of the culture by taking pictures every 0.5 minutes (**Fig. 1G**). We quantified activity as the number of pixels that changed intensity above background level between consecutive photos due to amoebae’s propagating behavior (**Fig. S2A**). Then, we expressed the cumulative sum of intensity change over time as *cumulative activity*. In all experiments, the cumulative activity over time displays an initial increase that is well described by a mono-exponential time course after a delay that reaches its plateau before 24 hours (**Fig. S2B**). This initial activity component reflects amoebae exiting the cyst state to become moving trophozoites that explore the substrate until they slow down to either feed on bacteria or become cysts again (**Movie S2)**. We termed the amplitude of this first plateau *cumulative inoculum activity (cIA,* **Fig. S2C**). For bacteria that support colonization by *Tetramitus*, a second increase in activity occurs due to amoebae behavior after feeding on the bacterial lawn: duplication, migration and encystation (**Fig. 1G**). These bacteria were *E. coli* OP50, *Comamonas, Stenotrophomonas* and the ensemble (**Fig. S2B**). We termed the second contribution to the cumulative activity plot *cumulative colonization activity* (c*CA*, **Fig. S2C**). The *ad-hoc* model described by equation 1 (Methods, **Fig. S2C**) was successfully used to fit experimental data and extract parametrized information, summarized in **Table 2**. Amoebae on bacteria that did not support colonization, *Chryseobacterium* and *Rhodococcus*, display similar activity curves than in unseeded NGM plates (**Fig. S2B**). This suggests that these two bacteria are either toxic or not eatable for *Tetramitus*, or both.

**Table 2.** Parameter details of amoebae tracking.

|  | NGM | *E. coli* OP50 | *Comamonas* | *Chryseobacterium* | *Rhodococcus* | *Stenotrophomonas* | ensemble |
| --- | --- | --- | --- | --- | --- | --- | --- |
| Cummulative inoculum activity, cIA<br>(A.U.) | 88.9 ± 70.4 | 60.3 ± 44.6 | 69.1 ± 26.6 | 95.2 ± 47.7 | 102.3 ± 53.2 | 149.8 ± 53.2 | 343.5 ± 190.8 |
| Relative colonization activity<br>(cCA/(cCA+cIA)) | 0.0 ± 0.0 | 1.0 ± 0.0 | 0.8 ± 0.1 | 0.0 ± 0.0 | 0.0 ± 0.0 | 0.9 ± 0.0 | 0.7 ± 0.1 |
| Excystation delay (tx, hr) | 4.6 ± 0.8 | 5.6 ± 1.8 | 3.5 ± 0.6 | 4.0 ± 0.2 | 2.6 ± 0.4 | 2.8 ± 0.2 | 2.6 ± 0.7 |
| Exploration time (tp, hr) | 9.3 ± 4.2 | 9.6 ± 2.3 | 9.0 ± 0.8 | 5.2 ± 0.6 | 9.6 ± 4.2 | 9.1 ± 2.1 | 5.5 ± 0.8 |
| Colonization time (tc, hr) | >200 | 52.7 ± 3.6 | 47.0 ± 2.9 | >200 | >200 | 67.7 ± 6.4 | 40.1 ± 12.2 |
| Growth time (tg, hr) | >200 | 2.0 ± 0.4 | 3.4 ± 0.3 | >200 | >200 | 1.8 ± 0.3 | 5.4 ± 1.3 |
| N | 3 | 3 | 4 | 3 | 4 | 3 | 3 |

Inoculum activities have similar features among all bacteria tested, displaying no statistically significant differences in amplitude (**Fig. S2D**), excystation time (*tx*, **Fig. S2E**) and exploration time (*tp*, **Fig. S2E**). This means that excystation and exploration behavior are not significantly altered by chemical cues released by tested bacteria.

In *E. coli* OP50 relative colonization activity (*cCA*/(*cCA+cIA*)) is larger (**Fig. 1H**) and faster (**Fig. 1I**) than in any other bacteria. Conversely, amoeba colonizes slowly the ensemble with also the smallest relative colonization activity (**Fig 1H-I** and **S2B**). These decreased parameters in the ensemble are likely due to the presence of *Chryseobacterium* and *R. qinshengii*, which do not support *Tetramitus* growth. These results show that *Tetramitus* has specific responses to the natural microbiota, and is likely a key player in shaping the microbial ecosystem.

### *C. elegans* feeds and develops in the ensemble

To introduce *C. elegans* in the ensemble we first tested their growth and other life history traits on monocultures and in the consortia with and without amoebae. We placed embryos on the bacterial lawns and quantified animals that reached adulthood at 72 hours. *C. elegans* growth was supported by all bacteria, but *Rhodococcus* and *Chryseobacterium* delayed their development compared to the rest of the strains. Notably, these are the two bacteria that do not support *Tetramitus* colonization. This retardation in development was however masked in both ensembles, with and without amoebae (**Fig. 2A**). Because nematode development is accelerated by vitamin B12 producing bacteria (38), we tested the production of B12 in monocultures and ensemble using nematodes expressing the *acdh-1* promoter driving *gfp* (39), which is turned off in the presence of the metabolite. *Comamonas*, the ensemble and surprisingly also *Rhodococcus* produce vitamin B12 (**Fig 2B**), suggesting that other metabolites or physiology of *Rhodococcus* may counteract the acceleration of development exerted by vitamin B12. Other consequences of feeding vitamin B12 producing bacteria to nematodes is the protection of neurons in a genetic model of degeneration where the Touch Receptor Neurons (TRNs) in *C. elegans* die by expression of the *mec-4d* degenerin (40). In the *mec-4d* paradigm, the AVM postembryonic neuron degenerates progressively in a stereotyped fashion after the birth of the animal (41). We recorded the morphology of the AVM neuron after feeding *mec-4d* animals with the ensemble and individual bacteria and classified the axons as functional or degenerated (**Fig 2C**). *Rhodococcus*, *Comamonas* and the ensembles provided neuroprotection with 20-30% of functional axons, coherent with their production of vitamin B12. *Stenotrophomonas* and *Chryseobacterium*, were similar to nonprotective *E. coli* OP50 (below 5%, **Fig. 2D**). This suggests that vitamin B12 is available to animals feeding on *Rhodococcus*, *Comamonas* and also the ensemble that contains both.

**Figure 2.**
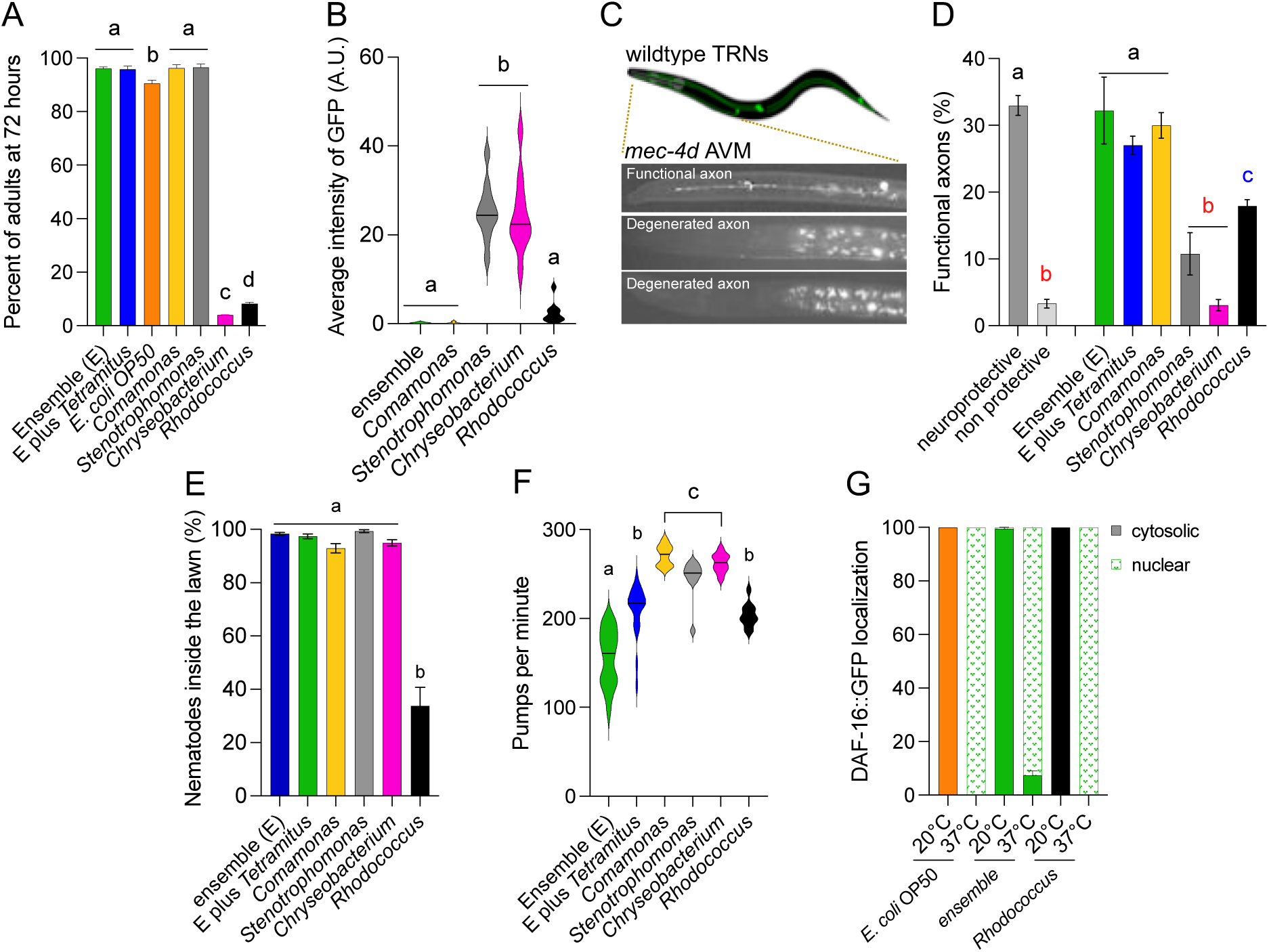
Introduction of *C. elegans* to the ensemble. **A.** Percentage of adult animals 72 hours after feeding embryos with the ensemble and monocultures. **B.** Arbitrary units (AU) representing average pixel intensity of *acdh-1::gfp* in animal intestine after feeding for 72 hours on the ensembles and monocultures. **C**. Scheme and images showing the touch receptor neurons (TRN) in wild type animals and the degeneration of the AVM (Anterior Ventral Microtubule) neuron in *mec-4d* animals. **D.** Percentage of functional AVM axons of animals feeding on the different bacterial conditions. **E**. Percentage of animals distributed in the areas with bacteria, either center or border after 72 hours in culture in different bacterial lawns. **F**. Quantification of pharyngeal pumping as pumps per minute of animals feeding on ensembles and monocultures. **G**. Percent of animals with cytosolic or nuclear localization of GFP driven by DAF-16 transcription factor, after being fed control bacteria, the ensemble and *Rhodococcus* at 20°C and at 37°C. Same letters denote there are no statistical differences, different letters mean statistical differences. Complete statistical analyses are detailed in **Dataset 2**.

Nematodes display dietary choices in bacterial lawns (42). We asked whether animals would locate in the center of the lawn, the border or outside, 72 hours after being placed outside but in close proximity to the lawns. In the ensemble with and without *Tetramitus* and in *Comamonas*, nematodes distributed between the lawn and an external border, which is made mainly of *Comamonas* growth, suggesting that animals prefer this bacterium over other in the ensemble (**Fig. 2E**). On *Stenotrophomonas* animals were on the lawn and mostly on an extended border, with less than 10% roaming on bacteria free spaces. *Chryseobacterium* caused animals to remain in the lawn. *Rhodococcus* was the only bacterium that induced animals to roam outside bacteria (**Fig. 2E**).

To study the feeding behavior of animals in the ensemble and monocultures, we measured pharyngeal pumping rate of nematodes living in the bacteria for a week. Ensembles with and without *Tetramitus* were not homogeneous lawns (**Fig. 1A**) and induced great variability in the pumping rates (**Fig. 2F**). Animals performed stops for a few seconds in one part of the lawn and then resumed at a high rate on a different domain of bacteria. Monocultures of *Comamonas*, *Stenotrophomonas* and *Chryseobacterium* induced the highest pumping rate while nematodes on *Rhodococcus* slowed pumping. The monocultures induced much more homogeneous responses, consistent with animals having a single food choice.

We observed that a number of animals feeding on monocultures of *Rhodococcus* showed the *Dar* (Deformed anal region) phenotype, previously observed with *Microbacterium nematophilum* (43), suggesting probable pathogenesis. To test whether *Rhodococcus* triggered a stress response we used a strain expressing a transgene for DAF-16 coupled to GFP (44). This strain shows nuclear localization of GFP upon starvation, thermic and pathogenic stress (21, 45). Animals were fed with *Rhodococcus*, the ensemble and the control bacteria *E. coli* OP50. **Fig. 2G** shows that DAF-16 is cytosolic at 20°C with all three bacteria, but a heat shock at 37°C induced nuclear localization in all conditions. This result suggests that neither *Rhodococcus* nor the ensembles are sensed as detrimental or pathogenic by the nematodes. Likely animals roam outside the *Rhodococcus* lawn because this bacterium is a large and hard to eat bacillus (46).

While we tested acute responses, we observed that culture plates of nematodes with the microbial ensemble lasted unusually long with reproductive animals and plentiful bacteria.

### Natural microbial ensembles are long lasting and induce nematode to diapause

To look closer into the long-term dynamics of the ensemble we co-cultured *C. elegans* with the microbial consortia at 20°C for eight weeks and monitored the growth of each species. We quantified bacteria as colony forming units (CFU), amoebae as individual cysts and nematodes by total numbers and developmental stage in one-week intervals. Initial co-cultures contained equal inoculums of each bacterium, a swab of amoebae and five wild type parental L4 *C. elegans* nematodes on each plate. Total bacterial CFUs in the ensemble remained constant after eight weeks (**Fig. 3A**), although individual bacterium fluctuated weekly (**Fig. 3A** and **Figs. S3A-D**). The organism counts after the first week of co-culture was considered the starting point to perform the comparisons with the following weeks. *Comamonas* was the only bacterium that decreased to 10^5^ CFU (**Fig. S3A**), suggesting *C. elegans* feeds on it more than on other bacteria*. Stenotrophomonas* peaked the second week driving a transient total CFU increase (**Fig. S3B**). *Chryseobacterium* CFUs decreased slightly throughout the eight weeks (**Fig. S3C**) while *Rhodococcus* remained constant (**Fig. S3D**). *Tetramitus* cysts were 10^8^ in the first week decreasing to 10^5^ by the eighth week (**Fig. 3A**) suggesting they are fed on by other organisms in the ensemble.

**Figure 3.**
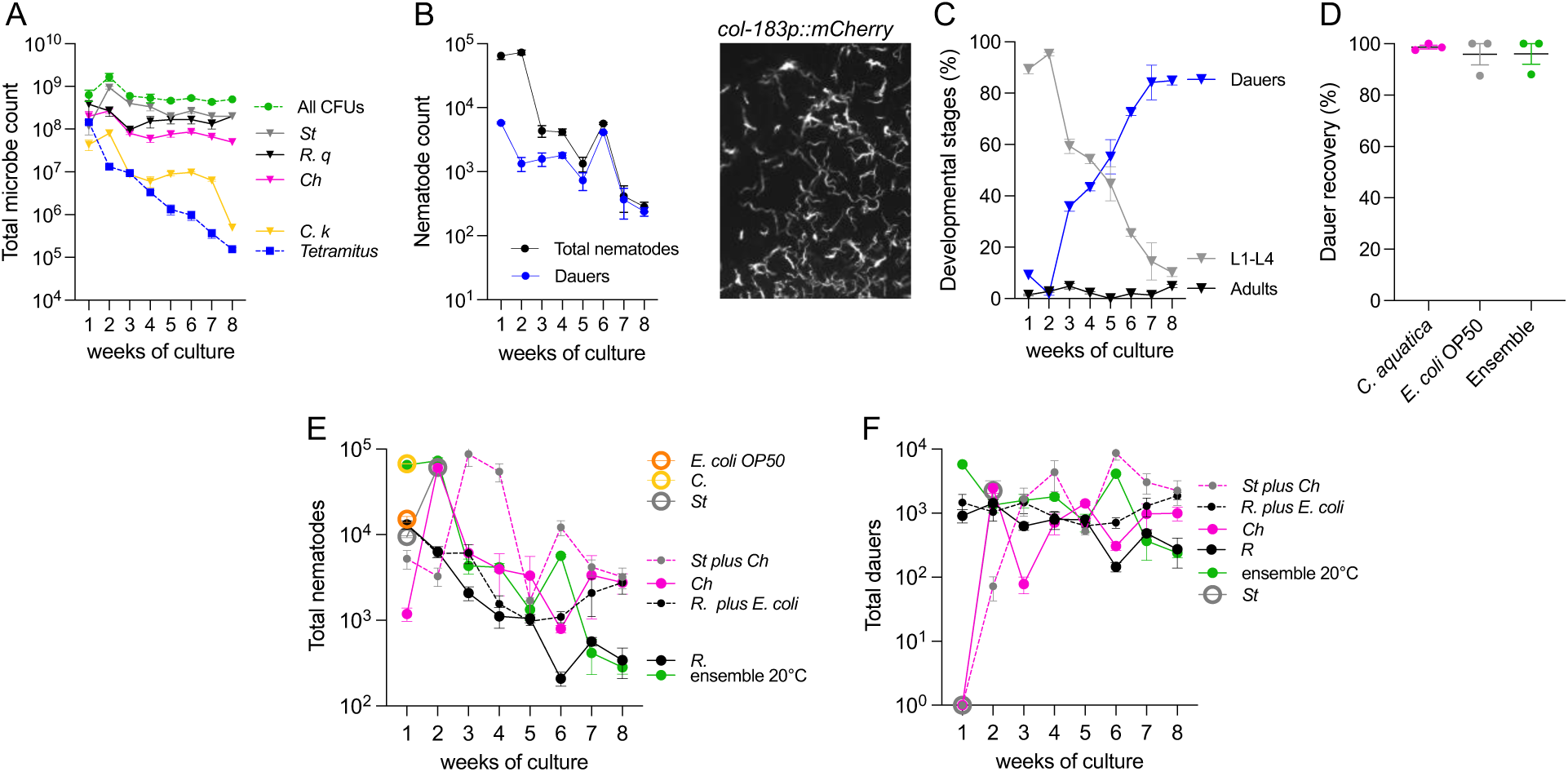
Characterization of population number of the components of the ensemble in a long-term co-culture experiment. **A**. Total microbial count, bacteria and amoeba in the ensemble in a long-term experiment at 20°C. Total bacterial CFUs are shown as well as the count of individual species in the consortia. **B**. Total nematodes and dauers in eight-week experiments. Inset photo shows red fluorescence of dauers changed to monochromatic, of the *col-183* promoter driving *mCherry*. **C**. Percentage of dauers, adults and non-adult nematodes in the ensemble in eight weeks. **D**. Percent of dauer recovery in three different bacteria. Animals that recovered were L4 at 24 hours and adults at 48 hours. **E-F**. Percent of total animals (**E**) and dauers (**F**) in the total population of nematodes grown for a week in different bacterial conditions. Complete statistical analyses for each graph are detailed in **Dataset 2**.

After one week there were 60×10^3^ total nematodes which decreased to three hundred by the 8^th^ week (**Fig. 3B**). Unexpectedly, a large number of F2 from initial L4 P0 nematodes entered diapause the first week under plentiful conditions (**Fig. 3B** and inset photo), showing dauers can be induced in the presence of non-pathogenic bacteria. Total numbers of dauers and other stages decreased toward the 8^th^ week (**Fig. 3B**; adults and larvae in **Fig. S3E**), but their percentage in the colony increased from 10 to 85% (**Fig. 3C**). We termed this response <u>Da</u>uer <u>F</u>ormation in <u>N</u>atural <u>E</u>nsembles or DaFNE. We asked whether long time dauers could exit diapause to colonize other bacteria. We collected two-week-old dauers from the ensemble with 1% SDS, a treatment that only dauers survive but also induces dauer recovery (8). We placed dauers in three separate conditions: *E. coli* OP50, *Comamonas aquatica* and ensemble lawns. 24 hours after, most animals recovered regardless of the bacteria they were placed in (**Fig. 3D**), and by 48 hours all of them were adults showing that dauers in the ensemble are viable and capable of colonizing other niches.

We tested whether monocultures were long-lived and if by themselves induced diapause. *Comamonas* like the control *E. coli* OP50 were exhausted in less than a week with an average of 67×10^3^ and 13×10^3^ animals respectively (**Fig. 3E**), all of which were non-dauer larvae (**Fig. S3H**). Monocultures of *Stenotrophomonas* had dauers in the second week (**Fig. 3F** and **S3H**) time by which bacteria were exhausted. This suggests that the persistence of *Comamonas* and *Stenotrophomonas* after eight weeks in the co-cultures is due to interactions with other components of the ensemble. *Chryseobacterium* and *Rhodococcus* monocultures however, lasted for eight weeks (**Fig. S3G and H**), with over 2000 and 300 total nematodes respectively at the end of the experiment (**Fig 3E**). *Rhodococcus* induced early dauers while *Chryseobacterium* produced dauers in the 2^nd^ week, which were maintained throughout the experiment (**Fig. 3F**). The co-culture of *E. coli* OP50 with *Rhodococcus* caused the former to last for 8 weeks (**Fig. S3I)**, and did not decrease the number of dauers in the culture (**Fig. 3F**). A similar effect was caused by *Chryseobacterium* to *Stenotrophomonas* when co-cultured, both lasting for 8 weeks (**Fig. S3J)**, and a dauer formation pattern that differed from either monoculture (**Fig. 3F**). We find that the ensemble is long lasting as are certain monocultures or combinations of bacteria. This shows that a diversity of environmental bacteria induces traits such as DaFNE that regulate nematode population growth.

### Nematode life history traits are specific of the ensemble and individual monoculture

The reduction of fertility could also contribute to avoid resource exhaustion. To assess aspects of the fertility and egg laying behaviors of nematodes in the ensembles, we counted the number of embryos laid by individual nematodes, the average amount of embryos *in utero* per animal, and their maturity in gravid *C. elegans* feeding on monocultures and the ensembles. To assess maturity, we used transgenic nematodes that express *gfp* under the *odr-1* promoter in the AWB and AWC, a marker of late embryogenesis (47) and **Fig. 4A**.

**Figure 4.**
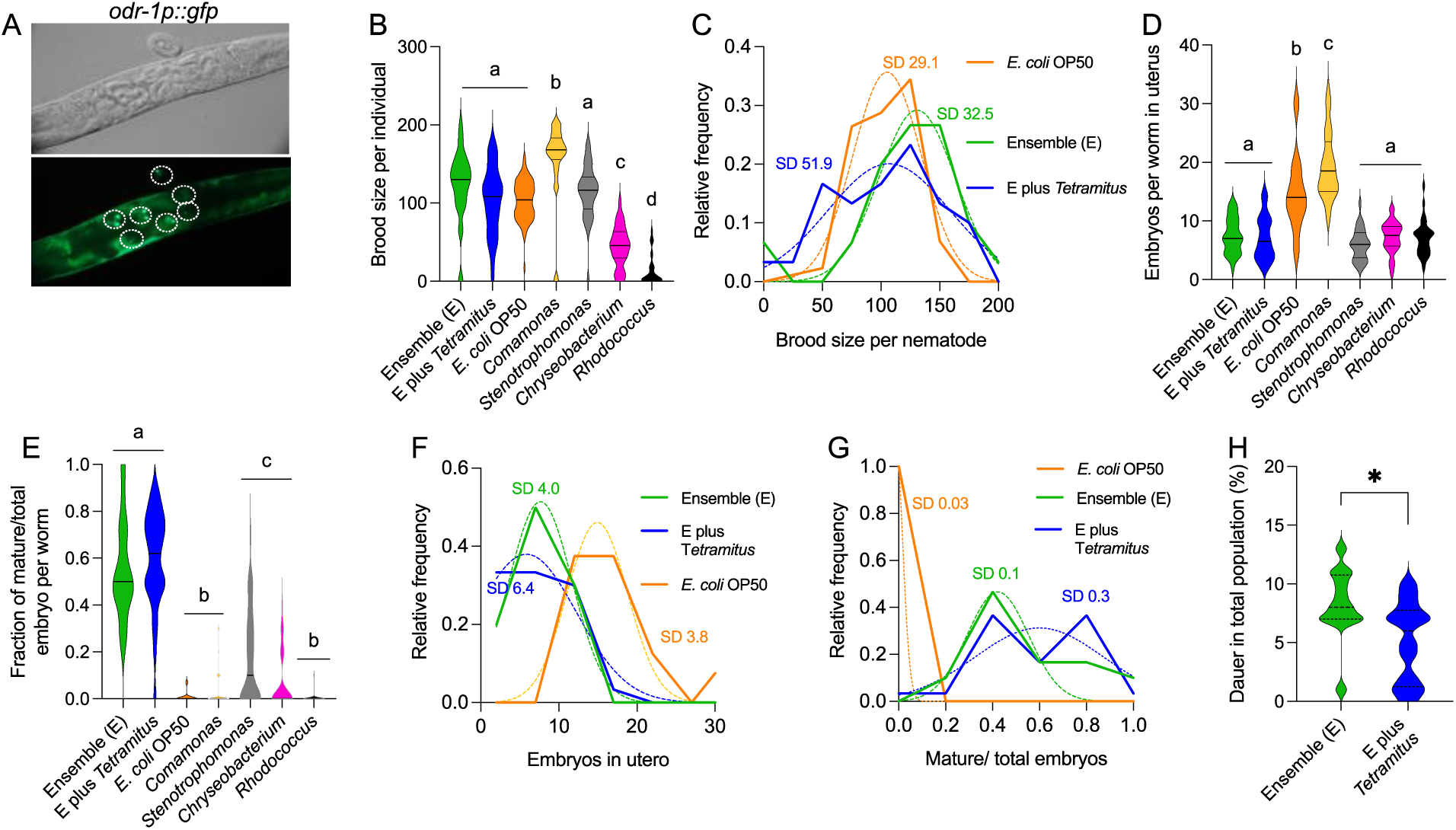
Fertility and maturity of embryos of nematodes growing on variations of the ensembles after one week in co-cultures. **A**. Nomarski (top) and fluorescence images (bottom) of nematodes expressing *gfp* in the AWB and AWC neurons under the *odr-1* promoter. White circles indicate individualized late-stage embryos as per their characteristic fluorescence pattern. **B-C**. Numbers of live progenies laid within a period of 72 hours by individual gravid adults (**B**) and the relative frequency of every experimental value within the population (**C**). The standard deviation for the condition of ensembles and ensembles with amoeba and the comparison with *E. coli* OP50 is shown. **D**. Number of embryos in utero after one week in the ensemble co-culture. **E**. Ratio of embryos expressing *gfp* in the AWB and AWC neurons vs total embryos in utero. **F-G**. Relative frequency of data points in the population of ensembles with and without amoebae compared to *E. coli* OP50 of **D** and **E**. **H**. Dauer percentage in animals feeding on ensembles with and without *Tetramitus*. * indicates pvalue<0.05 Same letters denote there are no statistical differences, different letters mean statistical differences. Complete statistical analyses are detailed in **Dataset 2**.

Nematodes on ensembles with and without *Tetramitus* had an average of 100 live progenies 72 hours after adulthood, not significantly different than *E. coli* OP50 (**Fig. 4B**). While the mean of those treatments is similar, the variation of the data is greater in ensembles with amoebae (**Fig. 4C**), suggesting that *Tetramitus* may affect the availability of certain species of bacteria in the co-cultures. Progenies *in utero* were far fewer in the ensembles (**Fig. 4D**) and the embryos were more mature than in *E. coli* or any other monoculture (**Fig. 4E**). Similar to the brood size, the variation of data points in the cultures containing amoebae was greater that on bacteria alone (**Fig. 4F** and **G**). To address the contribution of amoebae to DaFNE, we exposed the nematodes to ensembles with and without *Tetramitus*. While the trend in general is similar, the presence of amoebae lowers slightly the number of dauers in the population after one week (**Fig. 4H**).

Nematode brood size and embryos in uterus on monocultures of *Stenotrophomonas* were similar to the ensembles (**Fig. 4B**). The outputs in the nematodes in all other bacteria were significantly different to the consortia in most parameters (**Fig. 4B, D** and **E**). For example, *Comamonas* induced the largest brood size (**Fig. 4B**). Similarly, the number of embryos in uterus on animals feeding on *Comamonas* was high (**Fig. 4D**) with mostly immature progenies (**Fig. 4E**), suggesting that embryos are laid soon after they are made. On the other hand, *Rhodococcus*, though variable, caused worms to have very few progenies (**Fig. 4B**), which were laid in an immature state in the plates (**Fig. 4E**).

Because the consortium forms microdomains (**Fig. 1A**) it is likely that animals receive discontinuous contributions from bacteria and the amoebae that pray on them, that influence their life history traits.

### Bacteria and amoebae contribute differentially to the intestinal microbiota of *C. elegans*

We asked whether DaFNE required bacteria to be intact in the intestine. We grew cultures of the ensemble and used the bacteria-free supernatant and a lysate of the bacteria to supplement *E. coli* OP50 that were then fed to nematodes. Neither of these treatments caused DaFNE (**Fig. 5A**) showing that the communication between live bacteria and intestinal cells is needed for diapause induction. We then tested if on dauer-inducing conditions, bacteria from the ensemble were part of the intestinal microbiota of the nematodes. We quantified each species of intestinal bacteria by counting CFUs per nematode after one week in the ensemble. The predominant intestinal species was *Comamonas* (**Fig. 5B**) with an average 3×10^3^ CFU per nematode, followed by *Stenotrophomonas* and *Chryseobacterium* with 140 and 28 CFU respectively. *Rhodococcus* was very scarce, with an average of 7 CFUs (**Fig. 5B**), raising the possibility that this gram-positive bacterium cannot colonize well, or that animals choose other bacteria from the ensemble. We then isolated CFUs from animals fed exclusively with *Rhodococcus* and found consistently 10^2^ CFUs per nematode (**Fig. 5B**), ruling out an incapability of the bacterium to colonize.

**Figure 5.**
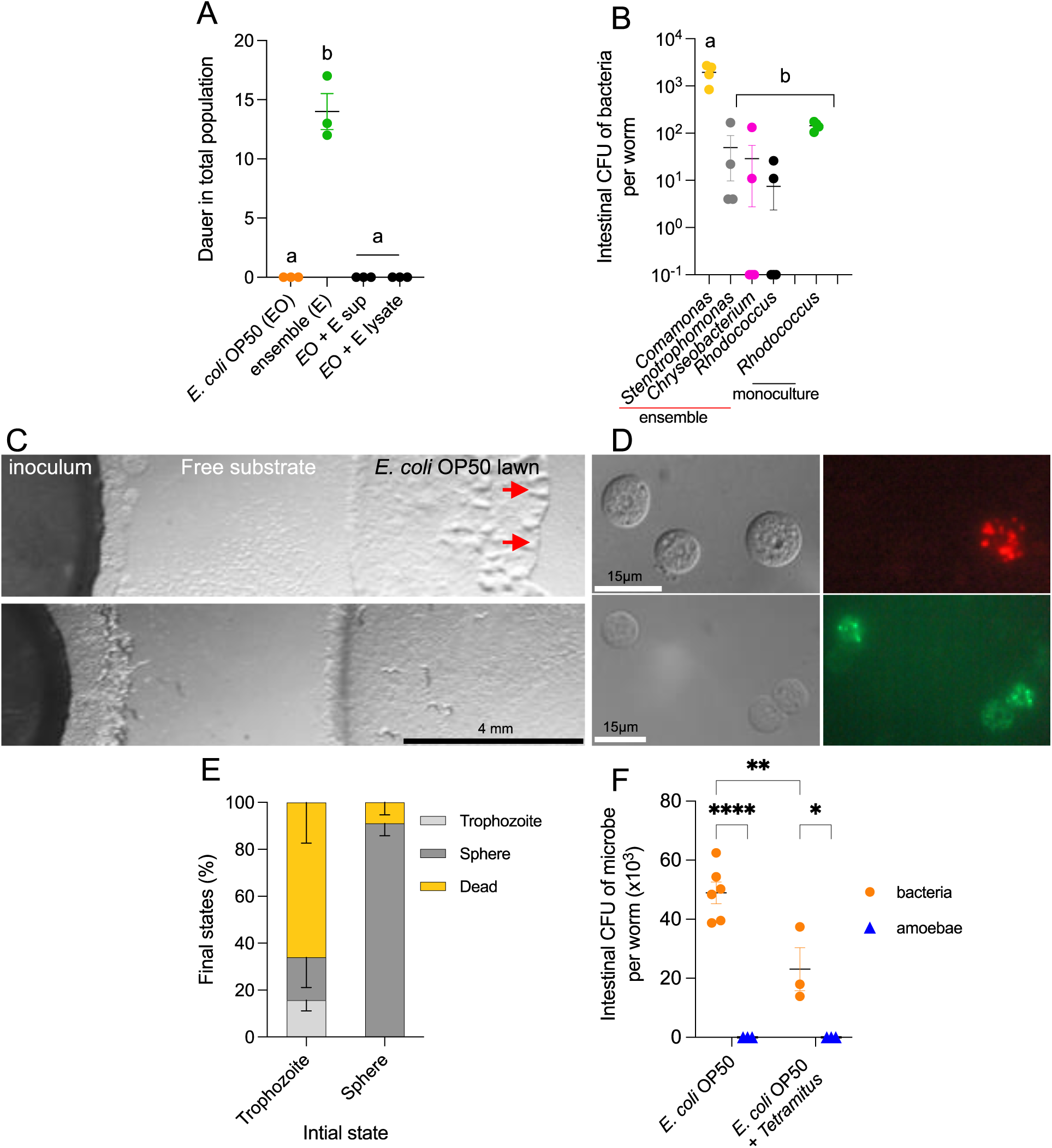
Behavioral parameters of nematodes feeding on ensembles with and without amoebae. **A**. Dauer quantification in cultures supplemented with ensemble’s supernatant and lysates. **B**. Intestinal CFU count of individual bacterium in animals feeding on the ensemble and monocultures of *Rhodococcus*. **C**. Colonization experiment of the ensemble containing amoebae into *E. coli* OP50-seeded NGM plates (top and bottom). Nematodes were placed only on the bottom experiment. The image was taken 3 days after placing synchronized L1s. A visible colonization front (red arrows) in the *E. coli* OP50 lawn is observed only in the plate without nematodes were amoebae cysts patches and individual *Tetramitus* are observed in the free NGM substrate. **D**. Cysts of *Tetramitus* after feeding on *E. coli* OP50 labeled with intracellular red and green fluorescent proteins. Note not all cysts show the fluorescent bacteria within their structure. **E**. Stages of amoeba with and without nematodes in a microscope pad observed at 20X. Sphere refers to either cysts or newly form round structures. **F**. CFU of intestinal bacteria or amoebae in nematodes fed with sole *E. coli* OP50 or *E. coli* and *Tetramitus* co-cultures. Same letters denote there are no statistical differences, different letters mean statistical differences. P value 0.1234 (ns); 0.0332 (*); 0.0021 (**); 0.0002 (***); 0.0001 (****). Complete statistical analyses are detailed in **Dataset 2**.

Another potential interacting microbe of *C. elegans* is the amoeba *Tetramitus. C. elegans* can feed on the amoebae *Dictyostelium discoideum* grown in axenic cultures (48). However, whether amoeba is a natural food of the nematode has not been studied nor has *C. elegans* been standardly grown on amoebae. *Tetramitus* feeds and multiplies rapidly on *E. coli* OP50 (**Fig. 5C** upper panel) and the bacterium can be found alive inside the amoebae cyst (**Fig. 5D**). When *C. elegans* is added to those cultures the growth of amoebae is restricted and the front of ameba growth is no longer observed (**Fig. 5C** bottom panel).

To directly observe whether nematodes are capable of feeding on *Tetramitus*, we mounted non-anesthetized animals with bacteria and amoebae on a microscope pad. We recorded the feeding behavior of *C. elegans* in the presence of motile and non-motile *Tetramitus*. *C. elegans* was able to feed on trophozoites but not the cyst form (**Movie S3,** quantification on **Fig. 5E**). In a number of cases, when pumping worms touch the amoebae, *Tetramitus* adopts a spherical shape in response to the suction exerted by the nematode (**Movie S4, Fig. 5E**).

We asked if amoebae could reproduce inside the animals by extracting the intestinal content of adult nematodes feeding on *Tetramitus*. We quantified the number of intestinal bacteria and amoebae of animals feeding on bacteria and *Tetramitus* and on bacteria alone after 72 hours. *Tetramitus* were not found inside the nematodes confirming they cannot survive to the sucking action when ingested (**Fig. 5F**). Second, animals that fed on bacteria alone had more intestinal CFUs than those feeding on bacteria together with amoebae, suggesting that nematodes that feed on *Tetramitus,* eat less bacteria.

### Temperature effect on ensemble components and diapause

Wild *C. elegans* have been found predominantly in mild to colder environments (49). We explored how increasing or decreasing by 5°C the temperature affects the growth of the ensemble and DaFNE in *C. elegans* in similar experiments to those shown in **Fig. 3**, done at 20°C. There were fewer animals at 15°C the first week compared to 20°C or 25°C, but this trend shifted by the third week with fluctuations that resulted in a larger population by the 8^th^ week (average 8×10^3^ nematodes, **Fig. 6A** and **S6A**). The first week at 25°C there were 4×10^3^ total animals, which decreased progressively to an average of 49 by the 8^th^ week (**Fig. 6A** and **S6B**). The emergence of dauers was slower at 15°C than at 25°C or 20°C, starting the second week (**Fig. 6A**), likely due to nematode quorum. At 15°C dauers were maintained at constant numbers between the 2^nd^ and the 8^th^ week (**Fig. 6A** and **S6C**). This was unique to the 15°C condition since both at 20°C and 25°C dauer numbers decreased progressively (**Fig. 6A** and **S6D**). Interestingly, at 25°C, DaFNE is triggered with less animals (average 4500 the first week) than at 20°C (65×10^3^) or 15°C (64×10^3^ the second week). The percentage of dauers in the total population of nematodes increased weekly, reaching 70% at 15°C and 94% at 25°C (**Fig. 6B**). The adult population fluctuated minimally at 15°C (**Fig. S6E**) while at 25°C the number of adults dropped throughout the weeks (**Fig. S6F**). Similarly, young larvae numbers (mostly L1) increased at 15°C (**Fig. S6G**) but diminished significantly at 25°C (**Fig. S6H**) by the 8^th^ week. This suggests that, at higher temperature, survival of the population depends on dauers.

**Figure 6.**
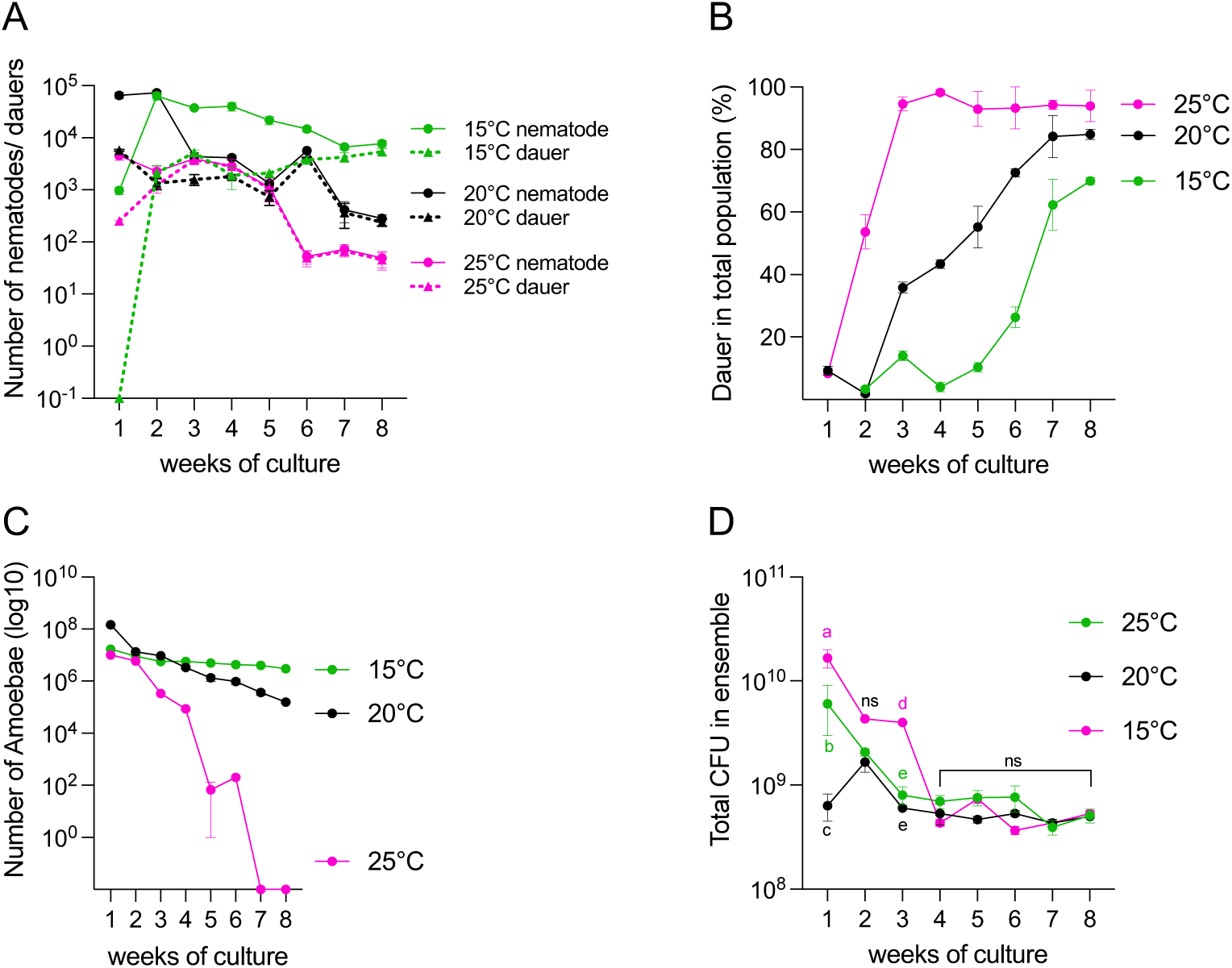
Effect of temperature in the ensemble dynamics. **A**. Total nematodes and dauers at each temperature. **B**. Percentage of dauers in the total population at each temperature. **C**. Number of individual cysts of amoebae (log10) in an 8-week temporal course. **D**. Total CFUs (sum of all bacterial CFUs) at each temperature each week. Statistical analysis comparing each temperature at each week is shown. Same letters denote there are no statistical differences, different letters mean statistical differences. P value 0.1234 (ns); 0.0332 (*); 0.0021 (**); 0.0002 (***); 0.0001 (****). Complete statistical analyses are detailed in **Dataset 2**.

Amoebae numbers at 15°C were maintained throughout the 8 weeks (approximately 3×10^6^), compared to 10^5^ at 20°C. At 25°C, *Tetramitus* decreased to exhaustion by the 7^th^ week (**Fig. 6C**), indicating that amoebae are not viable at sustained higher temperatures in the long term.

Bacterial CFUs were initially more numerous at 15°C than at 20°C or 25°C (1^st^ and 3^rd^ weeks), but after the 4^th^ week the total CFUs were the same at all temperatures (**Fig. 6D**). Individually, at 15°C *Comamonas* and *Stenotrophomonas* fluctuated the first three weeks (**Fig. S6I, J)**, while *Chryseobacterium* and *Rhodococcus* changed the 2^nd^ week and then remained constant (**Fig. S6K-L**). At 25°C, *Comamonas* and *Chryseobacterium* decreased the second week and then stabilized (**Fig S6I, K**) while *Stenotrophomonas* and *Rhodococcus* did not vary. This suggests that the ensemble self regulates to keep a similar quorum of total bacteria under the laboratory conditions at the three tested temperatures.

### DaFNE is a multigenerational response to natural bacteria

Crowding and the limitation of food are inductors of diapause (18). In the long-term paradigm, food is not exhausted and the percentage of dauers increase with the generations while the total nematode number decreases at all temperatures (**Fig. 7A-C**). This reflects that the relationship with quorum is influenced by other factors. For example, at 25°C the quorum needed to induce the first dauers is on average 5000 animals, almost 10 times lower that at 20°C or 15°C. To establish a baseline quorum for DaFNE we exposed different numbers of naïve P0 embryos to the bacterial consortium. After three days on the ensemble, over thirteen thousand naïve animals are needed to obtain 1% of dauers, and over twenty thousand for 10% (**Fig 7D**).

**Figure 7.**
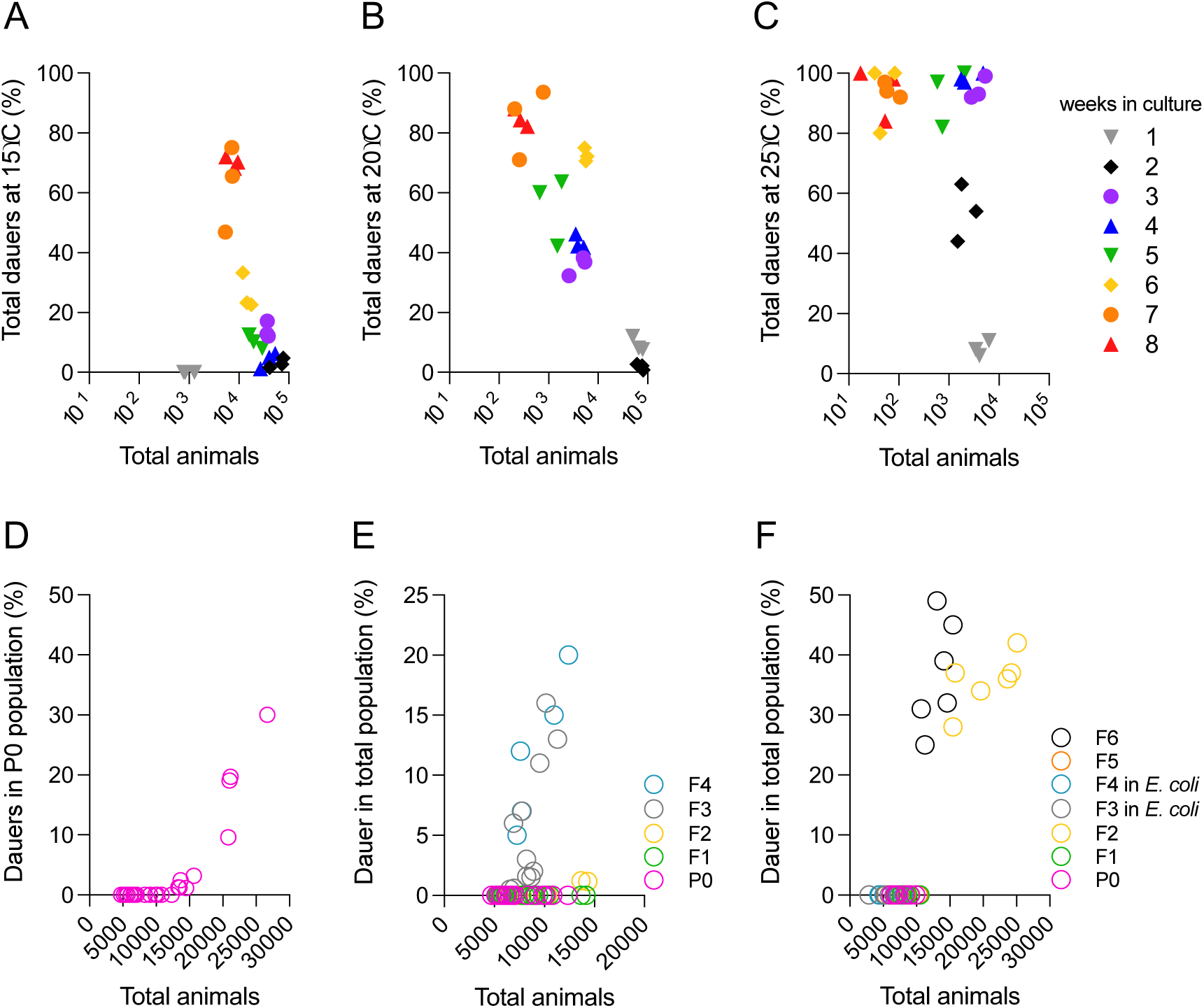
Effects of generational time on DaFNE. **A-C**. Percentage of dauers at 15°C (**A**), 20°C (**B**) and 25°C (**C**) in a long-lasting intergenerational paradigm without changes of bacteria or animals between weeks. **D**. Percentage of dauers in the P0 generation with increasing total numbers of animals. **E-F**. Percentage of dauers in an Intergenerational (**E**) and transgenerational (**F**) paradigm with interruption of each generation by hypochlorite treatment. Statistical analyses are detailed in **Dataset 2**.

Next, to quantify the multigenerational cumulative effect of the interaction with bacteria, each generation was extracted by hypochlorite treatment and the embryos were re-exposed to newly seeded plates of the ensemble. We defined that a multigenerational effect could be either inter or transgenerational (50). Intergenerational effects are those that accumulate as generations pass in the presence of the stimulus while transgenerational effects require the interruption for at least two generations of the dauer-inducing stimulus. To assess if DaFNE has an intergenerational effect, we started with P0 animals and measured the number of dauers in the F1-F4 descendants. In every generation we used under fifteen thousand embryos (below the threshold for dauer formation in P0 and F1). The F2 needed around 15000 nematodes to induce over 1% dauers similar to the P0s, while the F3 and F4 had many more dauers with half the quorum (**Fig. 7E**), in support of an intergenerational effect of DaFNE.

A transgenerational effect needs an enhanced (higher percentage of dauers) or faster DaFNE (a generation earlier) after a second encounter with the ensembles following two generations in standard *E. coli* OP50. While the magnitude of DaFNE is similar in the F6 and the F2, after the re-exposure dauers formed one generation sooner (**Fig 7F**) occurring in the F6 instead of the F7. Additionally, the quorum required to form dauers in the F6 was smaller than in quorum-induced diapause in the P0 animals (**Fig 7D**). These findings show that DaFNE accumulates intergenerationally and causes a transgenerational memory.

### Nematode pheromone synthesis and the RNAi machinery are needed for DaFNE

Animal quorum is sensed by a number of receptors that perceive pheromones (51, 52). We tested whether DaFNE needed pheromone synthesis by using *daf-22* mutants, deficient in one of the key enzymes for ascaroside production. *daf-22* animals do not form dauers on crowding (51) (53) or upon pathogenesis (21) but can diapause on high temperatures (20). *daf-22* mutants did not form dauers after one week on the ensemble (**Fig. 8A**) or after continuous exposure for eight weeks (**Fig. 8B**) showing that the pheromone is essential in this paradigm.

**Figure 8.**
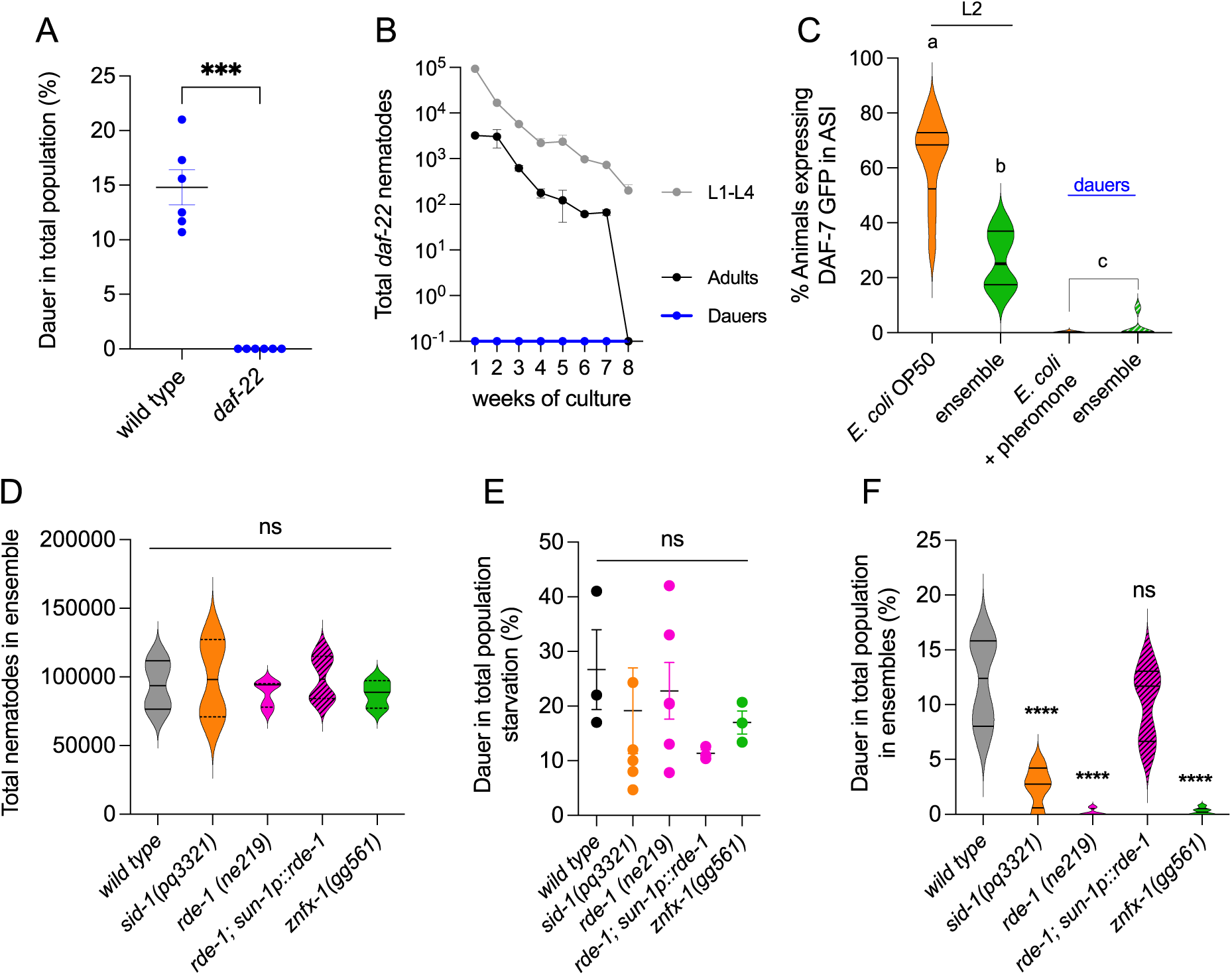
Mutations in the pheromone and RNAi pathways affect DaFNE. **A-B**. Percentage of dauer formation on wild type and *daf-22* mutants for one week (**A**) and total nematodes by developmental stage in a long-term paradigm of eight weeks (**B**). **C.** Percentage of GFP expressing animals in the ASI neuron under the DAF-7 promoter under varied conditions of bacterial food and development. **D-F.** Growth (**D**) and percentage of dauer formation in the population (**E-F**) of wild type, *sid-1*, *rde-1* and *znfx-1*mutant animals under starvation (**E**) and after a week on the ensembles (**F**). P value 0.1234 (ns); 0.0332 (*); 0.0021 (**); 0.0002 (***); 0.0001 (****). Complete statistical analyses are detailed in **Dataset 2**.

Pheromones suppress the expression of *daf-7,* the TGF-β (transforming growth factor-β) receptor ligand in the ASI neurons, causing dauer entry, while food restores both *daf-7* transcription and animal development (54, 55). We quantified the number of animals expressing GFP under the *daf-7* promoter in ASI neurons in the natural ensemble as developing animals and dauers. As reported, under standard feeding conditions, most L2 animals showed high expression of *daf-7* in the ASI neurons (63%), while most pheromone-induced dauers lost the expression (**Fig 8C**). In the ensemble, significantly fewer animals expressed high levels of *daf-7* (26%) compared to *E. coli* OP50 (**Fig 8C**). This suggests that the ensemble by itself reduces the expression of *daf-7* in the ASI compared to *E. coli*. Most dauers in the ensemble downregulated *daf-7* much like the pheromone (**Fig 8C**). This suggests that dauer induced by natural bacteria may have commonalities, although not identical, to the effects of pheromone.

Other molecular pathway involved in bacteria-nematode interspecies communication is the RNA interference (RNAi) machinery of the host and small RNAs from bacteria and host (12, 56–58). SID-1 mediates import of double stranded RNA (dsRNA) to cells (59, 60), and it is expressed in all tissues except the nervous system (61). Mutations in *sid-1* render RNAi originated in the intestine by bacterial production of dsRNA, ineffective (61). Cell-autonomous processing of this exogenous dsRNAs require the Argonaut RDE-1 (33, 34). Downstream events promote germline transmission of RNAi effects to the offspring (27), and for certain silencing RNAs in the germline, direct transgenerational epigenetic inheritance (TEI). TEI requires ZNFX-1, an RNA binding protein with zinc finger and helicase domains expressed uniquely in the germline (31). Because DaFNE is initiated by bacteria and is transmitted multigenerationally, we used mutants to test the role of the specific RNAi effectors mentioned above in dauer formation. We fed the ensembles to *sid-1 (pk3321), znfx-1(gg561), rde-1* (*ne219*) and *rde-1* (*ne219*) rescued in the germline (*sun-1p:: rde-1*) for one week and quantified their growth and ability diapause.

All mutants tested grew at wild type rates, reaching the quorum needed for DaFNE (**Fig. 8D**) and were capable of making dauers under starvation (**Fig. 8E**). DaFNE was impaired in *rde-1* and *znfx-1* mutants and diminished in *sid-1* animals (**Fig. 8F**). This suggest that the RNAi machinery is implicated in DaFNE at various levels: transport and RNA processing. Additionally, the germline rescue in *rde-1* mutants as well as its loss in *znfx-1(gg561),* highlights the relevance of germline transmission of RNAs for DaFNE. In summary, the response to natural bacteria involve cell to cell and autonomous communication based on RNAs and an ulterior epigenetic effect.

## Discussion

Hibernation and diapause are strategies used by many organisms to survive harsh environments. We report here that in natural long-lasting microbial ensembles of *Comamonas*, *Stenotrophomonas*, *Chryseobacterium*, *Rhodococcus*, and the amoeba *Tetramitus*, a portion of the *C. elegans* population enters diapause. We call this response Dauer Formation on Natural Ensembles (DaFNE). We speculate this can be a rationing strategy used under non-stressful conditions, more common than previously thought. It occurs in a wide range of optimal temperatures, from 15°C to 25°C; needs the nematode pheromone biosynthesis pathway and increases as generations pass and exposure persists, with a strong inter and transgenerational effect. In coherence with other multigenerational effects of bacteria, effectors of the RNAi interference machinery are required for DaFNE.

The amoebae *Tetramitus* displays a specific growth and feeding pattern on each bacterium of the ensemble. *C. elegans* in turn, also eats the trophozoites of the amoebae *Tetramitus* even when bacteria are plentiful. *Tetramitus* introduces variability to brood size, feeding behavior, and DaFNE among other traits. While the contribution of individual microbes to some traits can be notorious (i.e. *Rhodococcus* for dauer formation), our results show that the ensemble is a unit with distinctive traits that is not recapitulated entirely by an individual bacterium. The ensemble detailed here does not exhaust in experiments covering more than 16 *C. elegans* generations, offering a frame to study long lasting relationships in nature.

Nematodes have evolved superb strategies to colonize virtually every niche (4) (62) (63), pointing at a remarkable developmental and behavioral plasticity. One of these adaptations is their ability to diapause for long periods as dauer larvae, which is well suited for dispersal (64) (65) (66). The pioneer organism *C. elegans* has been instrumental in our understanding of the mechanisms of dauer formation (67) (68) (69), maintenance and recovery (70) mostly in the laboratory.

The ability of *C. elegans* to diapause offers significant benefits in populations that experience thermic stress (20), food shortage and overcrowding (18) or pathogenic infection (21). Even though these are stresses present in all environments, it is not known which are the cues regulating dauer entry in nature (22, 23). Interestingly, *C. elegans* dauers have been found in decaying fruits with sufficient bacteria of diverse species (10), suggesting that diapause may occur as a way of food rationing or in other words to avoid futile food consumption (71). It is unknown whether soil bacteria can induce dauer formation as a mechanism to induce fitness in the population by regulating population growth, in the absence of a canonical stress signal. In that sense DaFNE seem to be different than Pathogen Induced Diapause Formation (PIDF, (56), where animals diapause as a protective strategy against pathogenic infection. PIDF is also remembered for many generations after the stress is no longer present (12, 21). DaFNE is sustained for many weeks in the ensemble, and can recover in different bacteria including the ensemble itself. Remains to be explored whether consequences in the long run after DaFNE take place. For examples, long term diapause after feeding standard *E. coli* OP50, may cause negative tradeoffs to the immediate offspring but confers fitness to their F3 progenies increasing their resistant to starvation (72). Additionally, artificially selected dauers have progenies more prone to form dauers when stressed (73). Other effect of the bacterial ensemble is a strong egg retention in uterus and consequent matricide. Similarly, genetic variation in natural isolates of *C. elegans* in the egg laying circuit result in different degrees of egg retention (74). While this decreases maternal fitness, it increases offspring protection to chemical stress and faster development (74).

DaFNE has a cumulative effect as generations pass that does not depend on quorum. This multigenerational effect manifest in an intergenerational increase of dauers and also as an accelerated appearance of dauers after re-exposure. Thus far, transgenerational memory to bacteria in *C. elegans* have been observed using human or natural pathogens (21, 26, 58), and involves a stress response. Quiescence in natural environments in response to natural nonpathogenic bacteria may be a response to either control reproduction and thus preserve resources (71), or a mechanism to maintain a population resistant to sudden changes in abiotic factors or pathogenesis. For example, an abiotic accelerator of DaFNE is the increase in temperature in the optimal range, which also decreases the required quorum. Similarly, in one generation experiments, a rise in temperature also enhances pheromone induced diapause (75).

As in pheromone-induced dauers, DaFNE involves the repression of DAF-7/TGF-β expression in ASI neurons. DaFNE likely involves the sensing of bacterial molecules inside and outside the intestine as well as pheromones produced by community nematodes through olfactory neurons. Neuronal communication to the germline of social environment affects generational time in progenies of animals exposed to pheromones (52). Such neuron to germline communication may underlie the rescue of DaFNE by enabling *rde-1* Argonaute in the germline alone.

An open question is the nature of the bacterial metabolite that induces diapause. Given the dependence on the RNAi machinery, it is probable that a small RNA (sRNA) mediates this response as observed with *P. aeruginosa* PAO1 *RsmY* sRNA (12) in PIDF or *P. aeruginosa* PA14 *P11* sRNA in behavioral avoidance (26). Other type of microbial metabolites may also participate in the induction of DaFNE, as they are known to contribute significantly to the modulation of their host’s behavior (13–16). While some of these traits can be induced by the isolated metabolite (38, 40), relationships that causes adaptive behavioral responses require the active communication between microbe and the animal’s intestine (12, 21). This entity, called the holobiont is finally the unit capable of adaptation or adaptability (5, 76). In the present work, the abundance of each microbial species forming the *C. elegans*-ensemble holobiont, fluctuate in the intestine and in the culture plate. For example, *Comamonas* a vitamin B12-producing microbe is always dominant in the intestine, similar to the beneficial commensal *Pantoea*, selected by nematodes over other microbes in natural communities (7). *Rodococcus,* which causes a reduction in the progeny and delays development, is either absent or in small numbers in the intestine. Reasons for this could include the preference for other bacteria or the intestinal dynamics of the ensemble (77). *Rhodococcus*, unlike the strains considered as microbiome of *C. elegans* (78), is a gram-positive bacterium, suggesting the worm selects out hard to eat or large bacteria.

A novel component of the *C. elegans* interacting microbiota is the amoebae *Tetramitus*, which also displays defensive behaviors toward the nematode and specific feeding behaviors in the monocultures and the ensemble. A distant relative of *Tetramitus, Acantamoeba castellani* can change the virulence of *P. aeruginosa* in co-cultures, lowering its pathogenic potential in the nematode (79). This suggests the communication amoeba-bacteria impacts bacterial physiology, which in turn affects behavioral outputs in the nematode. *C. elegans* can feed on *Tetramitus* even in the presence of eatable bacteria, and *Tetramitus* responds specifically to the microbiota. In summary, bacteria, nematode and amoebae interact with one another to produce a long term self-sustained niche in the laboratory where pheromones, bacterial metabolites and the RNA interference machinery interact to induce animal quiescence.

## Supplemental Material

**Figure S1.**
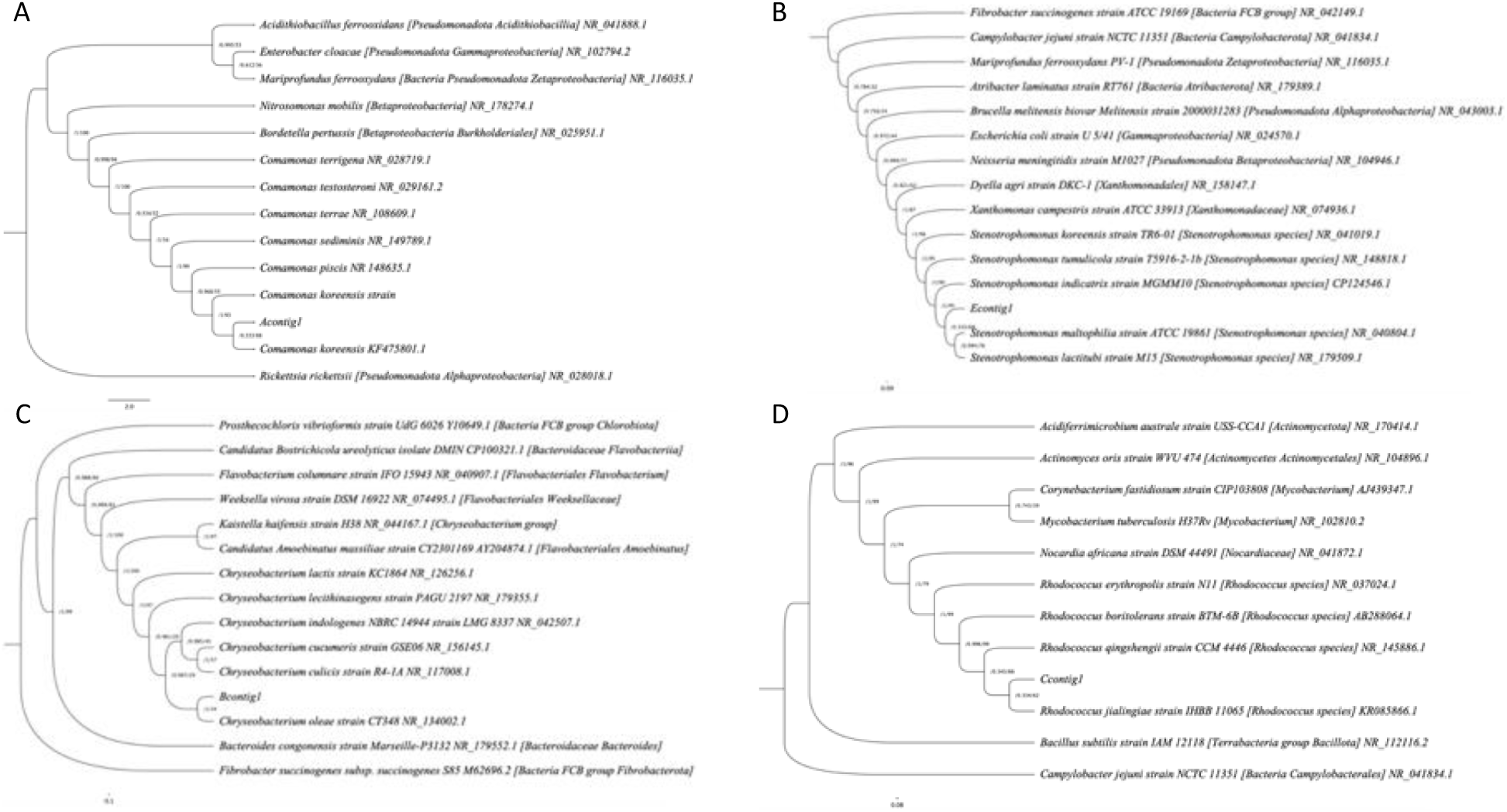
**A-D** Maximum Likelihood Phylogenetic tree of A-D contigs and surrounding taxonomic representative species. Bootstrap and Bayesian bootstrap values are displayed in the nodes.

**Figure S2.**
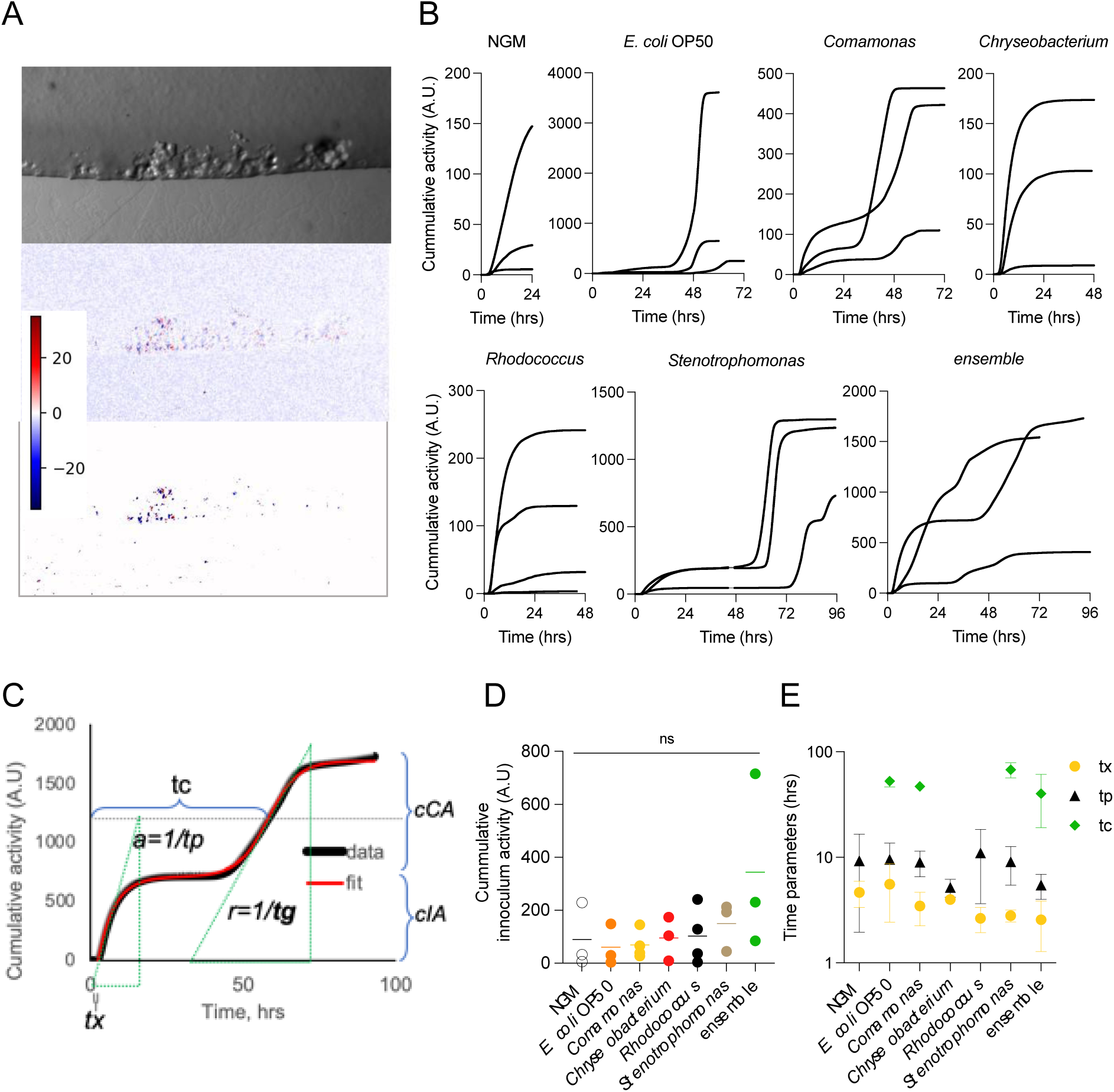
**A.** *E. coli* OP50-seeded NGM plate being colonized by *Tetramitus*. Top: a single picture of an *E. coli* OP50-seeded NGM plate where the bacterial lawn appears in the upper part of the photograph. Cysts were placed right below the bottom of the picture. After excystation, amoebae traveled the lower half to reach the bacterial lawn boundary, where they settle down and multiply (lumps near the agar/lawn interface). Note the trails left by moving amoebae connecting the bottom with the equator of the picture. Middle: pixel to pixel subtraction of two consecutive pictures taken 0.5 minutes apart. Bottom: pixel differences with absolute value smaller than 8 A.U. were discarded Activity due to amoebae moving around appear as intense blue and red spots. Intensity color scale is on the right (arbitrary units). Bigger patches of activity coincide with lumps in bacterial lawn near the center. **B.** Cumulative activity obtained from time lapse experiments performed on different bacterial lawns. NGM condition contains no bacterial lawn. **C.** *Tetramitus* colonization curve (black thick line) is an example of growth in *Stenotrophomonas* done by plotting the cumulative time-lapse activity over time. Red line is the model fit (see methods) that yields six parameters for each curve: tx, tp, tc, tg, cIA and cCA. **D.** Cummulative inoculum activity amplitude (cIA) for different bacteria. **E.** Excystation time (tx), Exploration time (tp) and colonization time (tc) obtained from parametric fit to equation 1. Experiments that do not support colonization (NGM, *Rhodococcus* and *Chryseobacterium*) have a colonization time tending to infinity (>>100 hours). A.U is arbitrary units. P value 0.1234 (ns). Complete statistical analyses are detailed in **Dataset 2**.

**Figure S3.**
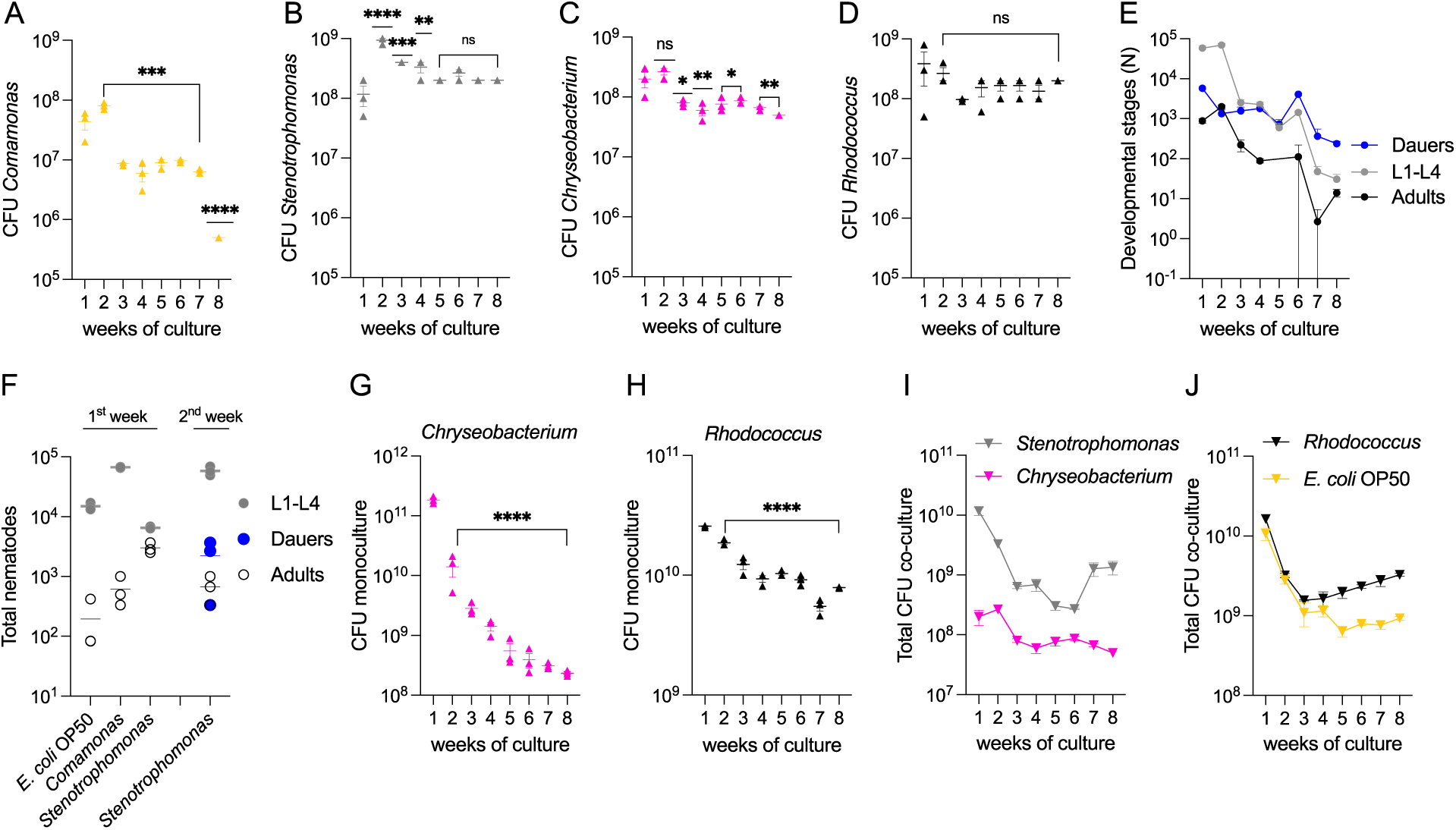
**A-D**. Number of individual CFUs per bacterial species in the ensemble (**A**) *Comamonas* (**B**) *Stenotrophomonas*, (**C**) *Chryseobacterium* and (**D**) *Rhodococcus* during the long-lasting co-culture. **E**. Progression of the numbers of adults, dauers and larvae in the ensemble throughout 8 weeks. **F**. Total nematodes as adults, dauers and larvae in monocultures of the control strain *E. coli* OP50 and *Comamonas* during the first week of growth and two weeks on *Stenotrophomonas*. **G-J**. CFUs of *Rhodococcus* (**G**) and *Chryseobacterium* (**H**) monocultures and *Rhodococcus* with *E. coli* OP50 (**I**) and *Stenotrophomonas* with *Chryseobacterium* (**J**) co-cultures during eight weeks growing with *C. elegans*. P value 0.1234 (ns); 0.0332 (*); 0.0021 (**); 0.0002 (***); 0.0001 (****). Complete statistical analyses are detailed in **Dataset 2**.

**Figure S6.**
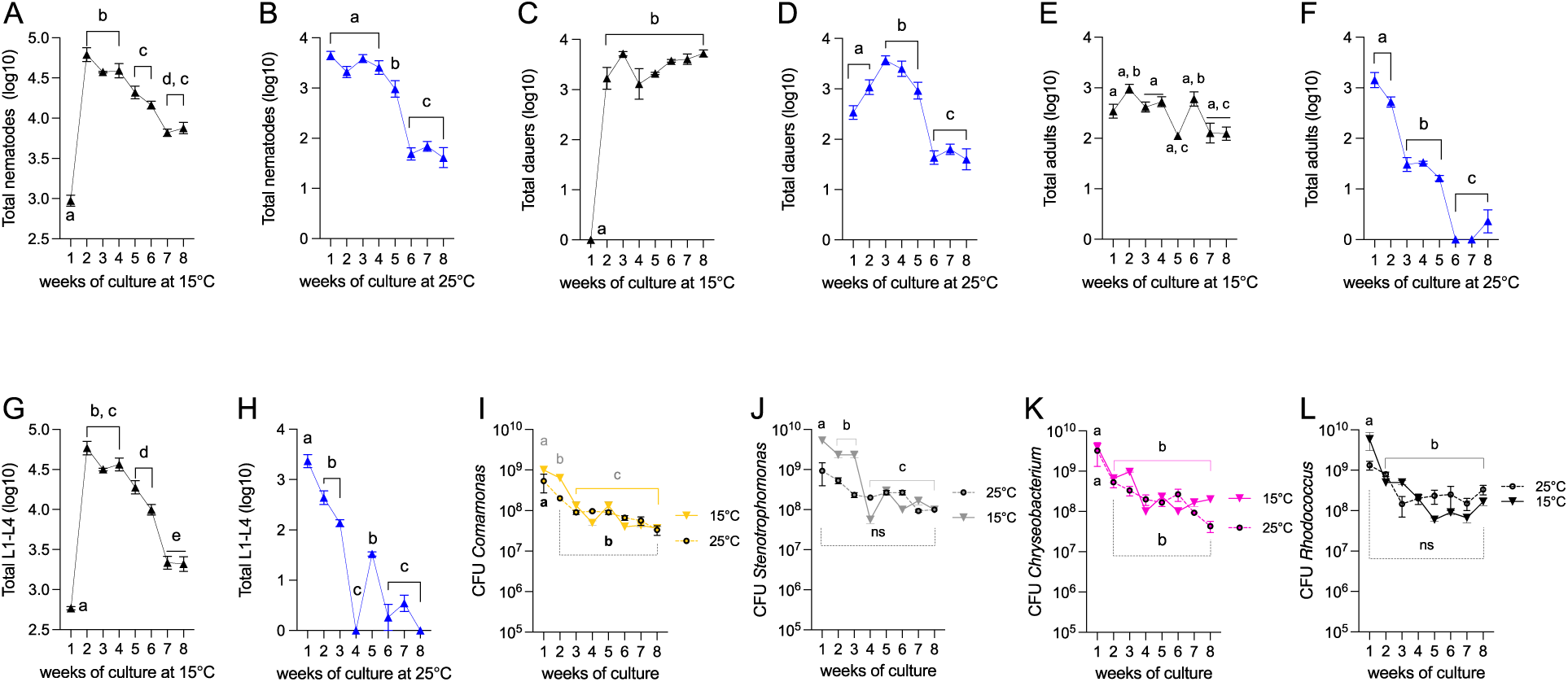
**A-H**. Total numbers of nematodes (**A, B**) and among those dauers (**C, D**), adults (**E, F**) and L1-L4 (**G, H**), in long term cultures at 15°C (**A, C, E, G**) and 25°C (**B, D, F, H)**. **I-L**. CFU count of *Comamonas* (I), *Stenotrophomonas* (J), *Chryseobacterium* (K) and *Rhodococcus* (L), in the ensemble at 15°C and 25°C. Same letters denote there are no statistical differences, different letters mean statistical differences. Complete statistical analyses are detailed in **Dataset 2**.

**Dataset 1.** Contains all the data in the manuscript.

**Dataset 2.** Contains all the statistical analyses in the manuscript.

### Materials and Methods

#### Formula for solutions and media used

M9 solution: 6g/L Na_2_HPO_4_; 3g/L KH_2_PO_4_; 5g/L NaCl; 1M MgSO_4_.

LB liquid media: 10g/L tryptone; 5g/L yeast extract; 10g/L NaCl, pH 6.8

LB solid media: 10g/L tryptone; 5g/L yeast extract; 10g/L NaCl, 15g/L agar; pH 6.8.

NGM media: 2% agar plates (0.25% Tryptone (2.5 g/L), 0.3% Sodium Chloride (NaCl) (3.0 g/L), 1.5% Agar, 1 mM Calcium Chloride, 1 mM Magnesium Sulfate, 25 mM Potassium Phosphate (KPO4) pH 6.0, and 5 µg/mL Cholesterol.

Hypochlorite solution (made fresh): 14% of Bleach (Clorox), 10% liquid 10N NaOH and 76% distilled water.

##### Collection and culture of natural microbial ensembles

Microbial ensembles were collected from naturally occurring moss in the soil of Chilean urban area of Santiago city (−33.432480912271856, −70.6147650371416) during summertime. A piece of moss of around 27 cm^3^ was placed in a petri dish containing distilled water for 1-2 hours. After incubation motile nematodes, rotifers and protozoans were visible in the dissection microscope. A small sample of wild nematodes and soil debris was placed on *E. coli* OP50 seeded agar plate using a worm pick. Bacterial inoculum containing *Tetramitus* grew at room temperature, for several days. Wild nematodes did not survive. *Tetramitus* spread rapidly over the *E. coli* OP50 bacterial lawn, replacing the entire lawn in 48-72 hours with cysts after inoculation, while orange-colored bacterial ensemble grew more slowly and steadily; the difference in grow rate allowed easy separation of amoebae and bacteria. Room temperature (RT) fluctuated approximately between 15 to 25 °C during the 24-hour period.

##### Maintenance and growth of bacteria

Bacteria were grown overnight on Luria Bertani (LB) plates at RT (natural ensembles) and 37°C *(E. coli* OP50) from glycerol stocks. Next morning, a scoop of the inoculum was cultivated in liquid LB at 200 rpm at RT for 48 hours for individual bacteria of the ensemble and the consortia, and 37°C for 6 hours for *E. coli* OP50. 150 µL of the bacterial culture was seeded onto 60 mm NGM plates and allowed to dry overnight before use.

##### Maintenance and growth of C. elegans

Wild-type, transgenics, and mutant *C. elegans* were maintained at 20°C, as reported previously (80). The following nematode strains were used: wild type (N2); PS8438 [*syIs600* (*col-183p*::mCherry + *odr-1p*::GFP)]; [TU2773 [*uIs31(Pmec-17mec-17::gfp); mec-4d(e1611)*X]; NL3321 [*sid-1(pk3321)*]; WM27 [*rde-1* (*ne219*)], DCL569 *mkcSi13 [sun-1p::rde-1::sun-1 3’UTR + unc-119(+)]*, YY996 *[znfx-1(gg561)]* and TJ356 [zIs356 [*daf-16p::daf-16a/b*::GFP + *rol-6(su1006)*]; DR476 [*daf-22 (m130)*]; FK181 [*ksIs2* [*daf-7p*::GFP + *rol-6(su1006)*]. All animals were maintained in *Escherichia coli* OP50 at 20°C before using or feeding with other bacteria.

##### Growth and maintenance of Tetramitus

*Tetramitus* plates are maintained in Nematode Growth Media (NGM) seeded with bacterial lawn. 100-1000 cysts are taken from a fully colonized plate using a platinum loop 1-2 mm in diameter. Standard bacterivore food bacteria *E. coli* OP50 strain is used otherwise stated. Viable cysts can be obtained after months of storage at room temperature in sterile conditions, even from completely dried agar plates. We used 1 to 7-day old fully colonized plates for cyst inoculation.

##### DNA extraction and the identification of bacteria and amoebae in the ensemble

To identify bacteria, we first exhausted the cultures in LB plates until isolates were obtained. Individual bacteria on LB plates were sent to Macrogen for sequencing. PCR of 16S rRNA genes was performed using 27F and 1492R primers, and sequencing is conducted using 785F and 907R primers, which are the inter-primers, to identify bacteria.

##### Genus identification and phylogenetic trees of bacterial 16S RNA sequence samples

For each sample, its 16S RNA sequence was used as query for a NCBI’s blastn (81) (default megablast option (82) using NCBI’s nucleotide (nt) database (83, 84) followed by the download of the sequence similarity tree generated by an implemented Neighbor-Joining method, in Newick format. Inspection of the results and the similarity tree allowed the genus identification. Since the similarity tree is not a phylogenetic tree, and the number of branches with very similar sequences from the same genus is overwhelming in the blast results, a biological sampling of surrounding species at different taxonomic levels can broaden the phylogenetic background of the query species, allowing its location more clearly in the bacteria tree. This can be done by displaying in a phylogenetic tree all 16S RNA sequences from the species of the same genus as the samples that resulted from the blastn, together with additional 16S RNA sequences from each of the taxonomic levels of the genus, in each case. The criterion for selection of the species for each taxonomic level was, when possible, to take bacterial species which name is similar to or representative of the taxonomic level name. Then, a Maximum Likelihood (ML) phylogenetic tree was searched from these sequences, including the query in each case. The taxonomy levels of each sample were taken from the Taxonomy Database from NCBI (83). The 16S RNA sequences were downloaded from the NCBI’s nucleotide Database.

The ML tree is searched from a starting tree made from a multiple sequence alignment build using MAFFT (85). As most of the downloaded 16S rRNA sequences are described as “partial sequences” we used MAFFT’s most accurate algorithm, L-INS-I, using the “localpaired” option with 1,000 refinement iterations and the default fasta format output. The result was input to iqtree2 (86) for model inference (option -m TESTONLY), followed by several runs of iqtree2 for ML tree space search. Each iqtree2 run specified the best evolutionary model (option -m); the number of bootstrap iterations, a thousand, (-b 1000); a perturbation option (-perturb), (we run three searches for each following perturbation value: 0.3, 0.5, 0.7 and 0.9) and; we used the option --nstop 500, which is the number of additional tries the program does when an impossible tree is reached. Moreover, the-abayes option for Bayesian bootstrap was used. A total of 12 simultaneous tree space searches per sample were performed using up to 48 CPU threads, option -t AUTO resulted in the using between 1 and 3 threads per process, and up to 20GB of RAM, during around 4 hours in a computer with 4 Xeon 3.1GHz 128 CPU threads, 256GB of RAM and 40TB of hard disk. The iqtree output files for one example run, including the log, the iqtree report, the maximum likelihood tree, the likelihood distances, the bootstrap trees and the consensus tree are available as supplementary materials (Supplementary File X1-X4). The tree was visualized with Figtreev1.4.4 (http://tree.bio.ed.ac.uk/software/figtree/).

##### Genomic DNA extraction and the identification of amoebae

Genomic DNA from amoebae was extracted using the Quick-DNATM Miniprep Plus Kit (Zimo Research), specifically we used the protocol for Biological Fluids and cells with the following modifications: Initially the sample is incubated with 200µL BioFluid & Cell Buffer with 20µL of Proteinase K. Instead of incubating at 55°C for 10 minutes, we incubated overnight.

For whole genome sequencing we send 920 total ng of gDNA of amoebae eluted in Elution Buffer from the Quick-DNATM Miniprep Plus Kit.

Library construction and sequencing was done by Macrogen using a Truseq Nano DNA library in a NovaSeq 6000 Sequencer 150 pair end (150×2bp) 10Gb/sample.

##### Phylogenetic Inference Amoebae

We started by listing *Tetramitus* relative species taken from previous inferred trees found in the literature (35, 87). The initial intention was to infer a phylogenetic tree using all orthologous genes that might be available among the related species. It turns out that only a handful of protist species related with *Tetramitus* have currently more than a few sequences publicly available in NCBI and secondary sources like PK10 (88). After downloading the gene sets from a few species for which their genomes are sequenced and inferring the orthologous groups using OMA (89), we realized there were not many orthologous genes shared among *Tetramitus* relative species in the currently available gene/genome information. The internal transcribed spacer, or ITS, from the 5.8S ribosomal RNA sequence is one of them, which is useful for species identification in different eukaryotic groups including fungi and plants (90, 91). Also, frequently available is the ribosomal RNA 18S partial sequence for a few relative protist species. Both are used for phylogenetic inference (37, 92). 18S rRNA sequence is one order of magnitude longer than the ITS, thousands vs. hundreds of nucleotides, and has been used in previous publications for *Tetramitus* phylogenetic inference (36, 93). We decided to generate trees using both sequences, separately and concatenated. Here, we show the concatenated alignment results as they are the same as “18S only” results. As we used both rRNA sequences we had to compromise to a smaller number of species that had both of them available, than the number of species that had only one of the two rRNA sequences.

Thus, we selected from NCBI’s Nucleotide Database (83,84) (http://www.ncbi.nlm.nih.gov/nucleotide/) the records of the longest ribosomal 18S and 5.8 rRNAs including the ITS1 and ITS2 sequences found, respectively, in the following species: *Allovahlkampfia* sp. (JQ271670.1 from PKD 2011b type strain PV66; LC106131.1); *Naegleria clarki* (JQ271691.1 from strain 2HZ; AJ566625.1 from type strain BG6); *Paravahlkampfia* sp. (GU230754.1 from A1PW2; AJ698857.2 from isolate 6/3Ab/1B); *Tetramitus dokdoensis* (KY463322.1; KY463323.1); *Tetramitus thermacidophilus* (AJ621575.1; AM950228.1 ecotype Pzc6); *Tetramitus thorntoni* (KT257696.1 from type strain SkGuaT; AJ698843.1 from type strain CCAP1588/8T); *Tetramitus jugosus* (KT257697.1 from type strain PrGuaT; AJ698844.1 from type strain CCAP1559/1T); *Tetramitus entericus* (KC164219.1 from clone CF1-11; AJ698856.1 from isolate C101); *Vahlkampfia avara* (JQ271723.1 from type strain 4171L; AJ698837.1 from type strain CCAP1588/1AT).

The sequences were gathered in a multiFasta file for each rRNA which was the input for MAFFT (85, 94) for multiple sequence alignment. As the downloaded rRNA sequences are described many times as “partial” we used one of MAFFT’s most accurate algorithms, L-INS-I, using the “localpaired” option with 16 default refinement iterations and the default fasta format output. Then, we input the concatenated, 18S and ITS multiple sequence alignments, to iqtree2 (95) for model inference (option -m TESTONLY), followed by several runs of iqtree2 for maximum likelihood tree space search. Each iqtree2 run specified the best model (-m TIM3+F+I+G4); the number of bootstrap iterations, a thousand, (-b 1000); a perturbation option (-perturb), (we run three searches for each perturbation value: 0.3, 0.5, 0.7 and 0.9) and; we used the option --nstop 500, which is the number of additional tries the program does when an impossible tree is reached. Finally, we also used the -abayes option for Bayesian bootstrap, although for communication purposes these values are not shown in the final figure. A total of 12 simultaneous tree space searches were performed using up to 60 CPU threads, option -t AUTO varied between 1 and 5 threads per process, and up to 20GB of RAM, during around 10 hours in a computer with 4 Xeon 3.1GHz 128 CPU threads, 256GB of RAM and 40TB of hard disk. The iqtree output files for one example run, including the log, the iqtree report, the maximum likelihood tree, the likelihood distances, the bootstrap trees and the consensus tree are available as supplementary materials (Supplementary File X1-X7). The tree was visualized with Figtreev1.4.4 (http://tree.bio.ed.ac.uk/software/figtree/).

##### Extended Results Identification of Tetramitus

Currently, there are 33 species under the genus *Tetramitus* at the NCBI’s Taxonomy Database, out of which 23 are not classified. From the 10 *Tetramitus* classified species left, only 5 of them count with its 18S and the 5.8 rRNA including ITS1 and ITS2 sequences. The evolutionary model inferred as the best model according to Bayesian Information Criterion was TIM3+F+I+G4. The final Maximum-likelihood and consensus trees using this model and three trees from the perturbations 0.3, 0.5, 0.7 and 0.9, resulted with the same topology and very similar bootstrap values (Supplementary Files). The final trees were generated from 789 informative positions that resulted from the Multiple Sequence Alignment.

##### Extended Results Identification of bacterial species in the ensemble

In all four cases, the identification using blastn with nt database was clear at the genus level. Acontig1: *Comamonas*; Bcontig1: *Chryseobacterium*; Ccontig1: *Rhodococcus*; Econtig1: *Stenotrophomonas*.

The taxonomy of each genus, taken from NCBI’s Taxonomy Database is:

Acontig1: Bacteria; Pseudomonadota; Betaproteobacteria; Burkholderiales; Comamonadaceae; *Comamonas* (TaxID 160825).

Bcontig1: Bacteria; FCB group; Bacteroidota/Chlorobiota group; Bacteroidota; Flavobacteriia; Flavobacteriales; Weeksellaceae; Chryseobacterium group (TaxID 59732).

Ccontig1: Bacteria; Terrabacteria group; Actinomycetota; Actinomycetes; Mycobacteriales; Nocardiaceae (TaxID 1827).

Econtig1: Bacteria; Pseudomonadota; Gammaproteobacteria; Xanthomonadales; Xanthomonadaceae (TaxID 40323).

The representative 16S RNA sequences from each taxonomic level for each bacterial sample are (the format used was, species, NCBI accession number and taxonomic level that represents between brackets):

Acontig1: *Comamonas piscis* strain CN1, NR_148635.1 [Comamonas species]; *Comamonas sediminis* strain S3, NR_149789.1 [Comamonas species]; *Comamonas terrae* strain A3-3, NR_108609.1 [Comamonas species]; *Comamonas testosteroni* strain KS 0043, NR_029161.2 [Comamonas species]; *Comamonas terrigena* strain IMI 359870, NR_028719.1 [Comamonas species]; KF475801.1 *Comamonas koreensis* strain IHB B 1392 [Comamonas species]; *Comamonas koreensis* strain DC5, OM250441.1 [Comamonas species]; *Bordetella pertussis* strain 18323, NR_025951.1 [Betaproteobacteria Burkholderiales]; *Nitrosomonas mobilis* strain Nc, NR_178274.1 2 [Betaproteobacteria]; *Acidithiobacillus ferrooxidans* strain ATCC 23270, NR_041888.1 [Pseudomonadota Acidithiobacillia]; *Rickettsia rickettsii* strain R, NR_028018.1 [Pseudomonadota Alphaproteobacteria]; *Enterobacter cloacae* strain ATCC 13047, NR_102794.2 [Pseudomonadota Gammaproteobacteria]; *Mariprofundus ferrooxydans* PV-1, NR_116035.1 1 [Pseudomonadota Zetaproteobacteria];

Bcontig1: *Chryseobacterium lecithinasegens* strain PAGU 2197, NR_179355.1 [Chryseobacterium species]; *Chryseobacterium oleae* strain CT348, NR_134002.1 [Chryseobacterium species]; *Chryseobacterium indologenes* NBRC 14944 strain LMG 8337, NR_042507.1 [Chryseobacterium species]; *Chryseobacterium cucumeris* strain GSE06, NR_156145.1 [Chryseobacterium species]; *Chryseobacterium culicis* strain R4-1A, NR_117008.1 [Chryseobacterium species]; *Chryseobacterium lactis* strain KC1864, NR_126256.1 [Chryseobacterium species]; *Kaistella haifensis* strain H38, NR_044167.1 [Chryseobacterium group]; *Weeksella virosa* strain DSM 16922, NR_074495.1 [Flavobacteriales Weeksellaceae]; *Flavobacterium columnare* strain IFO 15943, NR_040907.1 [Flavobacteriales Flavobacterium]; *Candidatus Amoebinatus massiliae* strain CY2301169, AY204874.1 [Flavobacteriales Amoebinatus]; *Candidatus Bostrichicola ureolyticus* isolate DMIN MAG, sequence taken from chromosome CP100321.1:c61187-59649 [Bacteroidaceae Flavobacteriia]; *Bacteroides congonensis* strain Marseille-P3132, NR_179552.1 [Bacteroidaceae Bacteroides]; *Fibrobacter succinogenes* subsp. succinogenes S85, M62696.2 [FCB group Fibrobacterota]; *Prosthecochloris vibrioformis* strain UdG 6026, Y10649.1 [FCB group Chlorobiota];

Ccontig1: *Rhodococcus boritolerans* strain BTM-6B, AB288064.1 [Rhodococcus species]; *Rhodococcus jialingiae* strain IHBB 11065, KR085866.1 [Rhodococcus species]; *Rhodococcus qingshengii* strain CCM 4446, NR_145886.1 [Rhodococcus species]; *Rhodococcus erythropolis* strain N11, NR_037024.1 [Rhodococcus species]; *Nocardia africana* strain DSM 44491, NR_041872.1 [Nocardiaceae]; *Corynebacterium fastidiosum*, strain CIP103808, AJ439347.1 [Mycobacterium]; *Mycobacterium tuberculosis* H37Rv, NR_102810.2 [Mycobacterium]; *Actinomyces oris* strain WVU 474, NR_104896.1 [Actinomycetes Actinomycetales]; *Acidiferrimicrobium australe* strain USS-CCA1, NR_170414.1 [Actinomycetota]; *Bacillus subtilis* strain IAM 12118, NR_112116.2 [Terrabacteria group Bacillota]; *Campylobacter jejuni* strain NCTC 11351, NR_041834.1 [Bacteria Campylobacterales];

Econtig1: *Stenotrophomonas koreensis* strain TR6-01, NR_041019.1 [Stenotrophomonas species]; *Stenotrophomonas indicatrix* strain MGMM10 sequence taken from chromosome CP124546.1:c3236787-3235243 [Stenotrophomonas species]; *Stenotrophomonas maltophilia* strain ATCC 19861, NR_040804.1 [Stenotrophomonas species]; *Stenotrophomonas lactitubi* strain M15, NR_179509.1 [Stenotrophomonas species]; *Stenotrophomonas tumulicola* strain T5916-2-1b, NR_148818.1 [Stenotrophomonas species]; *Xanthomonas campestris* strain ATCC 33913, NR_074936.1 [Xanthomonadaceae]; *Dyella agri* strain DKC-1, NR_158147.1 [Xanthomonadales]; *Escherichia coli* strain U 5/41, NR_024570.1 [Gammaproteobacteria]; *Brucella melitensis* biovar Melitensis strain 2000031283, NR_043003.1 [Pseudomonadota Alphaproteobacteria]; *Neisseria meningitidis* strain M1027, NR_104946.1 [Pseudomonadota Betaproteobacteria]; *Mariprofundus ferrooxydans* PV-1, NR_116035.1 [Pseudomonadota Zetaproteobacteria]; *Fibrobacter succinogenes* strain ATCC 19169, NR_042149.1 [Bacteria FCB group]; *Atribacter laminatus* strain RT761, NR_179389.1 [Bacteria Atribacterota]; *Campylobacter jejuni* strain NCTC 11351, NR_041834.1 [Bacteria Campylobacterota]; The best evolutionary model according to BIC, in each case, was:

Acontig1: TIM3+F+I+G4; Bcontig1: GTR+F+I+G4; Ccontig1: TIM3+F+G4, and Econtig1: TN+F+I+G4. In all cases, all final ML trees using all perturb values, 0.3, 0.5, 0.7 and 0.9 had the same topology with very similar support values.

##### 8-week experiments

###### Initial culture

Individual ensemble bacteria (*Comamonas, Stenotrophomonas, Chryseobacterium, and Rhodococcus*) were grown from a glycerol stock kept at −80°C, into LB solid plates (90 mm) overnight at room temperature (18-25°C). Amoebae are maintained on NGM plates seeded with *E. coli* OP50 at room temperature. Next morning, a scoop of each monoculture plus a scoop of amoebae were inoculated in 15 mL of liquid LB and shaken at 200 rpm for 48 hours at RT. From that culture, 150 µL are seeded into NGM plates (60 mm) and allowed to dry/ grow for 48 hours to allow amoebae growth on the ensemble. Next day 5 L4 nematodes of the strain PS8438 [*syIs600* (*col-183p::mCherry* + *odr-1p*::GFP)] are placed onto the 60 mm NGM plates (24 individual plates per experiment) and allowed to grow either at 15°C, 20°C or 25°C for eight weeks.

###### Quantification of ensemble

Every week for 8 total weeks, 3 plates were examined per condition to quantify each species of the ensemble. Entire plates are washed with 1mL of M9 collected into a 1.5 mL Eppendorf tube and centrifuged for 2 minutes at 376xg. The supernatant is collected in a clean Eppendorf tube for bacterial quantification. The pellet is used for nematode and amoebae quantification.

###### Bacteria

From the supernatant dilutions were made starting from 1:10 to 1:10^7^. 150 µL of each dilution were seeded on solid LB plates in triplicates. Bacteria were counted as colony forming units (CFU), 72 hours after inoculation.

###### Amoebae

Amoebae cysts were counted from the nematode pellet and supernatant containing bacteria. Total amoebae were the sum of both counts. Serial dilutions starting 1:10 to 10^3^ were used to count in three separate measurements the numbers of amoebae cysts present in either sample.

###### Nematodes

Nematode pellet was resuspended in 500 µL of M9. 10 µL or 1:10 dilutions were used to count: 1. Total nematodes, 2. dauer larvae, 3. Adults and 4. L1-L4 larvae. Dauers were counted under the dissecting fluorescent scope using the *col-183::mCherry* marker only expressed in diapausing animals. An average of three measurements per condition were done for every count.

##### Supplementation of E. coli with ensemble supernatant and lysate

Individual bacteria of the assembly were grown overnight on LB plates at room temperature from its glycerol stock at −80°C. The next morning, a “scoop” of each culture was grown in 20 mL of liquid LB at 200 rpm, room temperature for 48 hours. The culture was centrifuged in Beckman Coulter Allegra 21R Centrifuge with a Beckman S4180 Swing Bucket Rotor at 5450xg for 10 min to separate the pellet and supernatant from bacteria. The supernatant was filtered twice with syringe-driven filters (30 mm by 0.2 µm; Biofilter) 150 µL of the filtrate was taken and added to NGM plates seeded with *E. coli* OP50. 1 mL of Lysis Buffer from the DNeasy PowerSoil Pro Kit, Qiagen was added to the pellet and incubated for 30 min at 25°C (room temperature), centrifuged at 5450xg for 10 min, and 150 µL of the lysate supernatant was taken and added to NGM plates seeded with *E. coli* OP50. Five L4 (PS8438) nematodes were deposited on the plates. The number of dauer was quantified 7 days later using as negative control animals in *E. coli* OP50 and positive control animals feeding on the ensembles.

##### Quantification of dauer larvae

###### Using a fluorescent marker

To quantify dauers we used the PS8438 strain which expresses a *col-183::mCherry* marker, unique of dauer larvae. The entire nematode population on each plate was collected in 1 mL of M9 medium. Ten microliters of a 1:10 dilution was scored 3 times to count the total population and the amount of dauers under a Nikon SMZ1270 fluorescence dissection microscope. Animals with a clear dauer morphology and expressing mCherry in the cuticle were scored as dauers.

###### Using 1% SDS

Wild type or mutant animals are washed off the plates in 1 mL, the pellet is resuspended in 500 µl. To count the total population a 1:10 dilution is made onto M9. To count dauers, a 1:10 dilution is made onto 1%SDS (10 µl in 90 µl of 1%SDS). Animals are incubated for 15 minutes on the 1%SDS solution. 10 µL of each dilution were used to count live dauers and total animals in the population. Dauers were plotted as their percentage in the total population.

##### Quantification of multigenerational effects of DaFNE

###### Intergenerational paradigm

Individual bacteria of the ensemble were grown overnight on LB plates at room temperature from glycerol stocks (−80°C). Next morning, an inoculum of each culture was grown in liquid LB with 200 rpm agitation, at room temperature for 48 hours. 150 μL of the culture were plated on 60 mm NGM plates, and allowed to dry/grow for 48 hours before use. PS8438 embryos obtained from hypochlorite treatment were deposited on the plates. The numbers of eggs deposited depended on the quorum range evaluated. We used between 5,000 to 30,000 embryos. Hypochlorite treatment was carried out as follows: The plate containing gravid adults was washed with 1 mL of M9 and the content was placed in a 1.5 mL Eppendorf tube. The tube was centrifuged at 376xg for 2 minutes and the supernatant was removed. 500 μL of hypochlorite solution were added to each tube and incubated for up to 5 minutes until embryos were visible and hermaphrodite corpses were almost dissolved. Tubes were centrifuged at 376xg for 2 minutes, washed with 1 mL of M9, centrifuged again and the supernatant removed. Resulting eggs were counted and placed in the plates with the ensembles. 72 hours post hypochlorite treatment, the total nematodes and dauers were counted. Every new generation was obtained from hypochlorite treatment of the parental animals and the resulting embryos were placed on newly seeded plates.

The quantification of total worms and dauers is done as follows: Each plate is washed with 1 mL of M9, collected in a 1.5 mL Eppendorf tube, centrifuged at 376xg for 2 minutes, the supernatant is discarded and the nematode pellet is resuspended in 1 mL of M9. 10 μL of the nematode solution is deposited on a slide and the total number of nematodes and the number of dauers were counted using a Nikon SMZ1270 fluorescence dissection microscope. Dauers were counted as animals expressing the *col-183::mCherry* marker. After counting, the nematode pellet tubes are centrifuged, the supernatant is discarded and 500 μL of hypochlorite solution is added and the hypochlorite protocol is continued as described above until the F4.

###### Transgenerational paradigm

Bacteria and nematodes were grown and treated identically to the intergenerational protocol. What differed between the two paradigms was that F2 embryos were placed on *E. coli* OP50 plates for three days (F3 in *E. coli*) and the F3 embryos as well (F4 in *E. coli* OP50. Embryos from the F4 were then re-exposed to the ensembles (the F5) and their embryos were passed for another generation to the ensembles.

###### Dauer recovery

Dauer animals were separated from non-dauer nematodes by 1% SDS treatment. Pellets of dauers were placed on plates seeded with *E. coli* OP50, *C. aquatica* or the ensemble (*Comamonas, Stenotrophomonas, Chryseobacterium* and *Rhodococcus*), in a drop outside the bacteria. Dauer exit was scored at 24 hours by counting the L4 animals or non-dauers that moved toward the bacterial lawn or were roaming outside the lawn. At 48 hours the numbers of adults were scored as a further step to confirm exit from the dauer state. Additionally, plates were observed under the fluorescence dissecting scope to score the presence of dauers by expression of the *col-183p::mCherry* mark.

###### Pharyngeal pumping

Ten well fed adult *C. elegans* in each bacterial lawn or the ensemble were used for the quantification of the number of pumps of the pharynx in one minute. We used a manual counter to register each pump and stopped when 1 minute had passed using a timer. The animals were observed under 10X magnification in a Nikon SMZ745 dissecting scope.

###### Microscopy, photography and video

Microphotography of *Tetramitus*, *C. elegans* and natural bacteria cultures was performed at low magnification directly on NGM/agar plates using a Nikon SMZ 745T trinocular dissection microscope with additional 2x front lens coupled with a 10x tube lens to a Sony XCD-SX910 digital camera via firewire to a PC.

Micro-assemblies were also imaged at high magnification using DIC optics on either a Nikon Eclipse Ni or Ti, equipped with Canon EOS Rebel t3i and Nikon D780 digital cameras, respectively. Samples were loaded onto 2 % W/V agarose pads on a 2 mm glass slide and a 0.17 mm coverslip on top.

Photograph of PS8438 dauers [*syIs600* (*col-183p*::mCherry + *odr-1p*::GFP)] expressing cuticular mCherry were imaged directly on their growing plate in a Nikon fluorescence dissection microscope SMZ 1270 at 40x magnification and using an adequate filter cube for GFP imaging. The CMOS sensor Alvium 1800 U-500 (Allied Vision) was exposed for 220 ms at a gain of 24.1 db.

###### Criteria for AVM neuron integrity

Morphological evaluation of AVM neuron was modified from ref. 41. Neurons with full-length axons, as well as those with anterior processes that passed the point of bifurcation to the nerve ring, were classified as AxW. Axons with a process connected to the nerve ring were classified as AxL, and those that did not reach the bifurcation to the nerve ring were classified as AxT. Lack of axon, only soma, and soma with only the ventral projection were classified as Axϕ. Functional axons (AxF) are AxW and AxL. We plotted only the AxF axons.

###### Time course of neuronal degeneration

Embryos resulting from hypochlorite treatment were placed in plates at 20°C with the desired bacterial food. We scored the integrity of the AVM neuron axon at 72 hours post-bleaching. For each evaluation, we used at least three biological replicas with triplicates of 30 worms each.

###### Pheromone extraction

A pheromone mixture was obtained as described by Schroeder and Flatt (96) with the following modifications: a one-liter culture of PS8438 nematodes started from si× 10 cm plates with starved animals, was grown for one month at RT (18°C-25°C), agitated at 110 rpm.

Experimental NGM plates were made with a concentration of 5%v/v of pheromone and 2% w/v heat killed *E. coli* OP50 bacteria was seeded 24 hours prior to experiments. 15 FK181 adult hermaphrodites were manually handled with a pick to the pheromone-NGM plates for 3 hours, allowed them to lay eggs and then removed. Plates were in an incubator for 72 hours at 20°C. Dauer larvae were selected manually by morphology and scored. 50-100 L2s were grown from a bleach in NGM plates seeded with live *E. coli* OP50 24 hours prior to incubation and maintained at 20°C for 24 hours.

###### Tracking of amoebae and behavioral analysis

*Tetramitus* culture development was studied using time-lapse photography by taking pictures of bacterial plates freshly seeded with hundreds of cysts. A fire-polished glass Pasteur pipette was gently dipped in fully colonized of *Tetramitus*. Cysts were transferred by touching the agar next to the bacterial lawn with the cyst-loaded tip at 1-2 mm away from the lawn border. Photos were recorded every 30 seconds for 48-96 hrs, using 40x magnification setting using image acquisition tools and scripts in Matlab (Mathworks). For time-lapse experiments, activity quantification was assessed by subtracting consecutive images and taking the absolute value of intensity change for each pixel in the picture matrix. The total number of pixels where intensity change was above certain threshold was counted and reported as active area vs time. The threshold was defined by inactive regions of the picture at initial stages of the time-lapse experiment, where subtle variations in illumination define the background noise.

An ad-hoc mathematical model was used to parametrize the colonization behavior described by the cumulative time-lapse activity (CA) over time, described by equation 1:

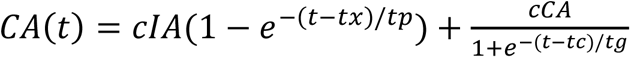

cIA and cCA are, respectively, the amplitudes of the inoculum and the colonization phases of the behavior. tx is related to excystation kinetics that defines the quiescence observed before a sudden rise in activity occur. tp is the natural decay time in exploration activity of moving trophozoites after excystation.

Phenomenologically, the mono-exponential relaxing inoculum component describes a first-order transition state of individual amoebae from fast moving (exploring) to very slow or settled, observed in all experiments.

tg (grow time) is the inverse of the natural grow rate of a logistic grow curve that describes the secondary colonization phase when present. tc (colonization time) is the time at which the cumulative activity of the colonization phase is half of its plateau and usually coincides with the moment of maximum instantaneous activity. Kinetics of amoebae colonization activity comprise cells feeding, growing in number and acquiring faster traveling speeds to subsequently exit the recording area and leaving behind immobile cysts in the original lawn area. The amplitude of the signal increases with larger number of amoebae moving in the field and also with the speed of individuals. However, we also found that the amplitude of the signal depends on the agar thickness and lawn coloring, giving rise to noncomparable illumination conditions between experiments. Therefore, we can only extract information from the relative amplitude of inoculum and colonization components.

###### Nematode brood size

Total brood size was counted for 30 individual F2 randomly picked adults grown for 1 week on bacterial natural ensembles at 20°C. The number of embryos laid and live larvae were at room temperature for three consecutive days, this is, re-counting new progeny each day to ensure total live progeny is being counted. For *E. coli* OP50, 90 individual adults were used.

###### Quantification of GFP expression in the ASI neurons

The strain FK181 [*ksIs2* [*daf-7p*::GFP + *rol-6(su1006)*] was used for quantification of GFP expression in the ASI neurons. Two different levels of expression were detected: Weak somas without neurite expression and strong fluorescence along the whole cell including soma and neurite. We plotted as GFP positive only the strong fluorescence. In both *E. coli* and ensemble conditions, dauers were isolated by 1% SDS treatment and confirmed by morphology. L2 animals were selected based on morphological criteria.

###### Quantification of GFP intensity in intestine

VL749 (*wwIs24 [acdh-1p::GFP + unc-119(+)]*) nematodes expressing intestinal *gfp* were imaged directly on their growing plate in a Nikon fluorescence dissection microscope SMZ 1270 at 40x magnification and using an adequate filter cube for GFP imaging. The CMOS sensor Alvium 1800 U-500 (Allied Vision) was exposed for 220 ms at a gain of 24.1 db. Pictures containing worms were analyzed using ImageJ by selecting the areas of the picture corresponding to individual animals and measuring average pixel intensity.

###### Bacterial count (CFU) in nematode intestine

We adapted the protocol from ref. 97 as follows: *Comamonas, Stenotrophomonas, Chryseobacterium, Rhodococcus* were inoculated on LB plates and cultured overnight at room temperature (18-25°C). The control *E. coli* was grown at 37°C. All bacteria are taken from glycerol stocks at −80°C. 24 hours later, an inoculum of each culture was mixed in liquid LB and agitated at 200 rpm. Ensemble plus amoebae is obtained by addition of a scoop of *Tetramitus* cysts. Ensembles were grown for 48 hours at RT and *E. coli* for 6 hours at 37°C. 150 µL of the cultures were plated on 60 mm NGM plates, and allowed to dry/grow for 48 hours before use. 5 L4 PS8438 animals were placed on triplicates with *E. coli* OP50, the ensemble and the ensemble plus amoebae *Tetramitus*. After 7 days 30 L4 worms of each condition were picked to a tube with 1 mL of M9. The tubes are centrifuged at 376xg for 2 minutes. The supernatant is then removed leaving up to 0.1 mL to avoid touching the nematode pellet. The pellet is washed with 1 mL of M9 supplemented with 25 mM levamisole (Lev+M9) and repeated six times. After this, 10 µL of the supernatant were used to seed an LB plate to control for microbes still remaining in the liquid with the nematodes after the washes (Control 1).

After this step, the tube containing the pellet was washed with 1 mL Lev+M9 plus a mix of Antibiotics [50 µg/mL Carbenicillin, 25 µg/mL Tetracycline, 25 µg/mL Chloramphenicol and 100 µg/mL Gentamicin] three times. After the last wash, the tube is incubated with Lev+M9+Antibiotics for 1 hour with agitation. During this step, after 30 minutes of incubation the solution is replaced with a fresh one for the remaining of the hour. Finally, the pellet is washed to eliminate the remaining antibiotics with 1mL Lev+M9 and centrifuged. After this, 10 µL of the supernatant were used to seed an LB plate to control for microbes still remaining in the liquid with the nematodes after the washes (Control 2).

The pellet is then washed again with 1 mL of Lev+M9 for three other cycles. After this, 10 µL of the supernatant were used to seed an LB plate as a final control of the protocol (Control 3). An extra control of 150 µL of the same supernatant is spread on an LB plate until exhaustion for colony quantification.

The worm pellet is then lysed with a pestle for 2 minutes or until they dissolve. The remaining lysate on the tube is filled with M9 up to 500 µL in the Eppendorf tube. Dilutions of 1/10 and 1/100 are made in M9 before placing 150 µL in LB in triplicates. Lysates containing *E. coli* OP50 are seeded on plates supplemented with 25 µg/mL of streptomycin. After incubation, the colonies are counted and the number of CFU/nematode is calculated according to ([CFU *dilution factor]/number of nematodes).

We used morphological macroscopic criteria to discriminate the individual bacteria. For example, *Comamonas* are small, white, clear, round colonies with a halo surrounding the colony, with a liquid texture. *Stenotrophomonas* are bigger than *Comamonas*, round, with a greenish color and viscous glossy texture. *Chryseobacterium* is bigger that *Comamonas* and *Stenotrophomonas*, round, strong yellow-orange color, extremely viscous. *Rhodococcus* is the smallest of all colonies, round, dark, with a non-homogeneous texture, with a pale pink matte surface.

### Biological and technical replicates and Statistical evaluation

Each experiment was performed in three technical triplicates and at least three biological replicates. All the data collected in this work is detailed in **Dataset 1**. We define biological replicates as experiments made on different days, containing triplicates of each condition, and a technical replicate as a triplicate of the same condition on the same day. Each figure contains at least three experiments (biological replicates) performed as explained before. All the biological replicates are performed spaced from each other from 1 day to 1 week.

Statistical evaluation was performed using one- or two-way ANOVA with post hoc tests and a Student t test when indicated. All statistical testing is detailed in **Dataset 2**.

## Acknowledgments

We thank Marcela Legüe for insightful comments on the manuscript. Rene Zúñiga and Carlos Vasconcellos helped with the codes for the time lapse acquisition. Some strains were provided by the CGC, which is funded by NIH Office of Research Infrastructure Programs (P40 OD010440). This work was funded by FONDECYT 122650 to AC.

## Author contribution

Conceptualization JPC and AC

Methodology MS, PO, GR, JPC and AC

Investigation MS, ER, GI, PO, GR, JPC and AC

Writing-Original Draft AC

Writing-Review and Editing PO, JPC and AC

Funding Acquisition AC

